# Mutation is a sufficient and robust predictor of genetic variation for mitotic spindle traits in *C. elegans*

**DOI:** 10.1101/033712

**Authors:** Reza Farhadifar, José Miguel Ponciano, Erik C. Andersen, Daniel J. Needleman, Charles F. Baer

## Abstract

Different types of phenotypic traits consistently exhibit different levels of genetic variation in natural populations. There are two potential explanations: either mutation produces genetic variation at different rates, or natural selection removes or promotes genetic variation at different rates. Whether mutation or selection is of greater general importance is a longstanding unresolved question in evolutionary genetics. Where the input of genetic variation by mutation differs between traits, it is usually uncertain whether the difference is the result of different mutational effects (“mutational robustness”) or different numbers of underlying loci (“mutational target”), although conventional wisdom favors the latter. We report mutational variances (VM) for 19 traits related to the first mitotic cell division in *C. elegans*, and compare them to the standing genetic variances (VG) for the same suite of traits in a worldwide collection *C. elegans*. Two robust conclusions emerge. First, the mutational process is highly repeatable: the correlation between VM in two independent sets of mutation accumulation lines is ~0.9. Second, VM for a trait is a very good predictor of VG for that trait: the correlation between VM and VG is ~0.9. This result is predicted for a population at mutation-selection balance; it is not predicted if balancing selection plays a primary role in maintaining genetic variation. Goodness-of-fit of the data to two simple models of mutation suggest that differences in mutational effects may be a more important cause of differences in VM between traits than differences in the size of the mutational target.

## Introduction

The question “What are the factors that govern genetic variation in natural populations?” has been central to the field of Evolutionary Genetics ever since its inception (Dobzhansky 1937; Lewontin 1974; Lewontin 1997). Within a group of organisms, seemingly similar or related phenotypic traits can vary considerably, and consistently, in the extent of genetic variation in the trait. For example, in many organisms, resistance to acute heat stress is much less heritable and evolvable than resistance to acute cold stress (Hoffmann *et al*. 2013). If different traits in the same set of organisms have consistently different levels of genetic variation, there are two potential underlying evolutionary causes: mutation and/or selection. Traits may differ in the mutational target they present, i.e., the number and/or types of loci that potentially affect the trait, or in the rate at which those loci mutate. Traits may also differ in the average effect that mutations have on the trait, i.e., they may be differently robust to the effects of mutation. Alternatively, traits may be subject to differing strengths and/or kinds of selection.

In the realm of quantitative genetics, a few empirical conclusions seem fairly certain. First, traits that are direct components of fitness – life history traits – are typically more genetically variable than other classes of traits (Houle 1992; Lynch *et al*. 1999). Second, life history traits experience greater input of genetic variation from mutation than other classes of traits (Houle *et al*. 1996; Halligan and Keightley 2009). Third, life history traits appear to be under stronger purifying selection than other classes of traits (Houle *et al*. 1996; Lynch *et al*. 1999; McGuigan *et al*. 2015).

A longstanding related question concerns the relative influence of balancing selection on the maintenance of genetic variation (Dobzhansky 1955; Lewontin 1974; Charlesworth 2015). Surely, a large fraction of mutations are unconditionally deleterious and are removed more or less efficiently by selection. However, if the mutation rate is high and selection is not too efficient, a large fraction of the genetic variation in a population may simply be deleterious junk loitering at mutation-selection balance (MSB). Alternatively, it is certainly possible that some alleles are subject to balancing selection, and even if mutations subject to balancing selection are rare, they may in aggregate explain a substantial fraction of the standing genetic variation (Barton 1990).

Houle (1998) investigated the relationship between the standing additive genetic variance (VA) and the per-generation input of genetic variation by mutation (the mutational variance, VM) for eight life history and morphological traits in *Drosophila melanogaster* and found a very strong positive association between VM and VA (Spearman’s *r* = 0.95, P<0.001; see Figure 1 in Houle (1998)). That result has a very clear interpretation: variation in mutation explains 90% of the variance in standing additive genetic variance *among traits*. Similar analysis of the genotypic variance (VG) and VM for nine morphological and life history traits in *Daphnia pulex* also reveals a strong positive correlation (*r* = 0.76, P<0.02; data in Tables 1 and 3 of Lynch *et al*. (1998)). A positive association between the mutational and standing genetic variance is predicted if most genetic variation is due to deleterious alleles at MSB (see below); it is not predicted by balancing selection, although it is theoretically possible (Houle *et al*. 1996; Charlesworth and Hughes 2000). However, Charlesworth (2015) recently analyzed five decades of quantitative genetic and molecular data from *D. melanogaster* and reached the conclusion that MSB cannot be a sufficient explanation for genetic variation for fitness in that species, hence there must be a significant contribution from balancing selection.

**Table 1.**
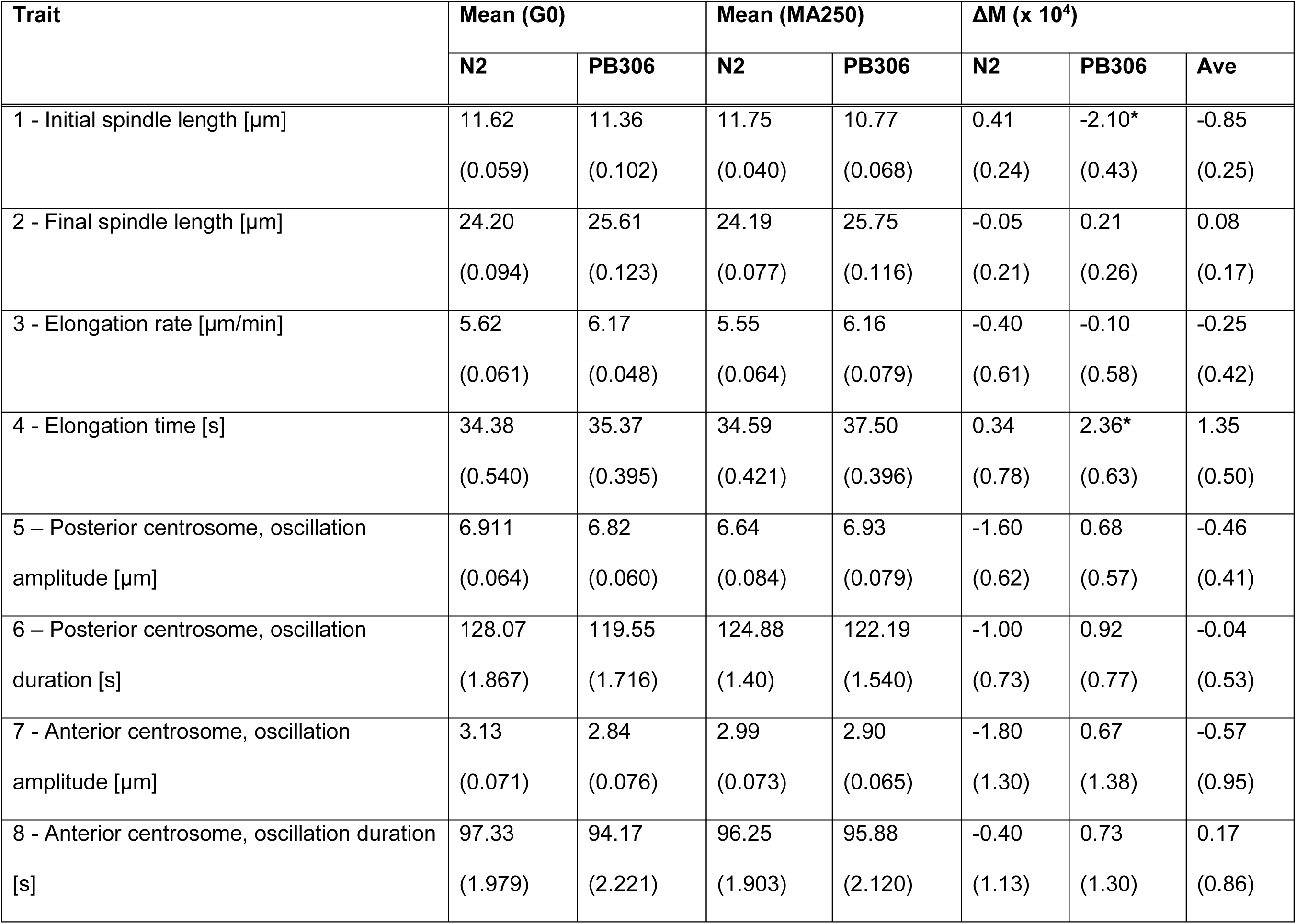
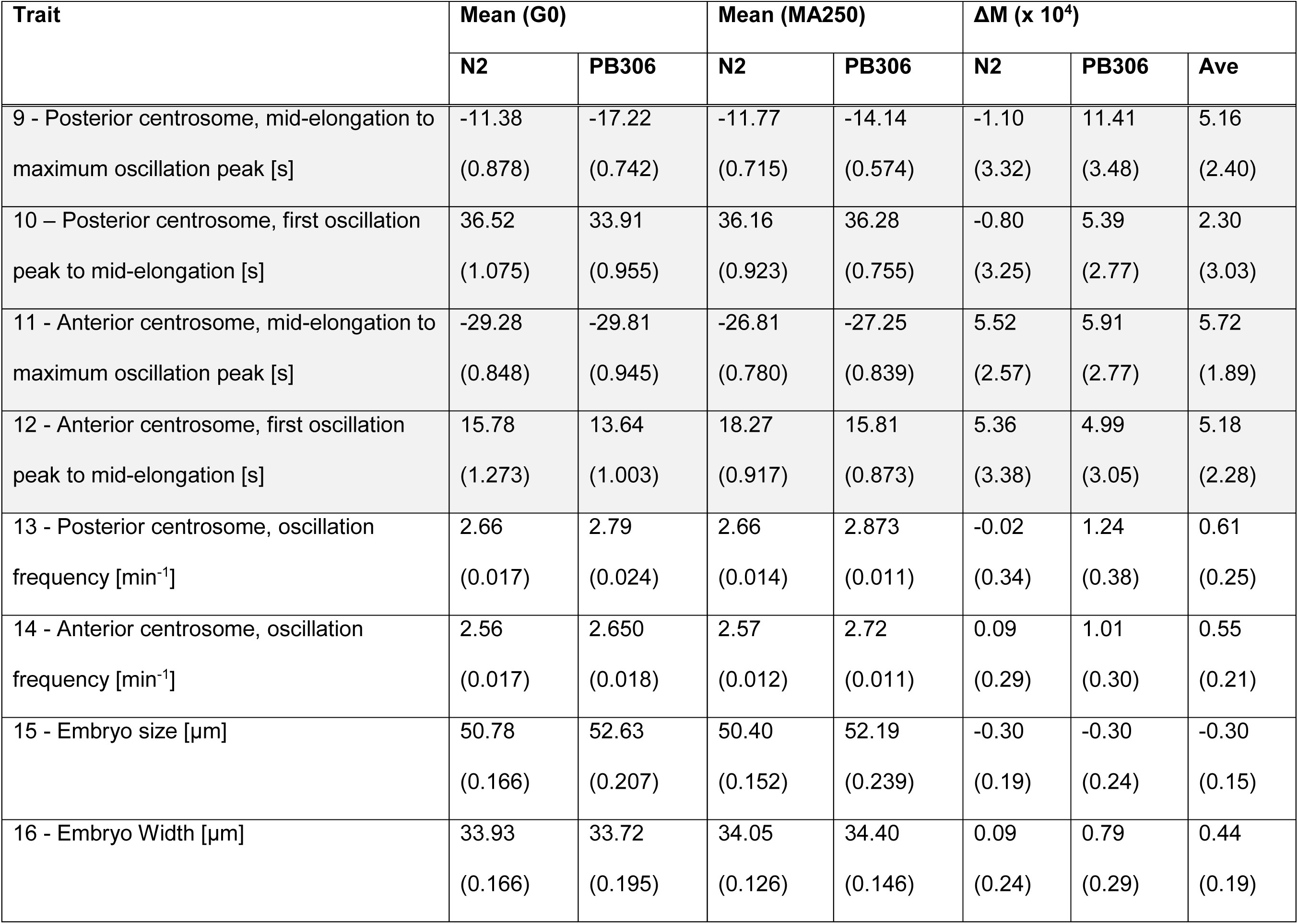
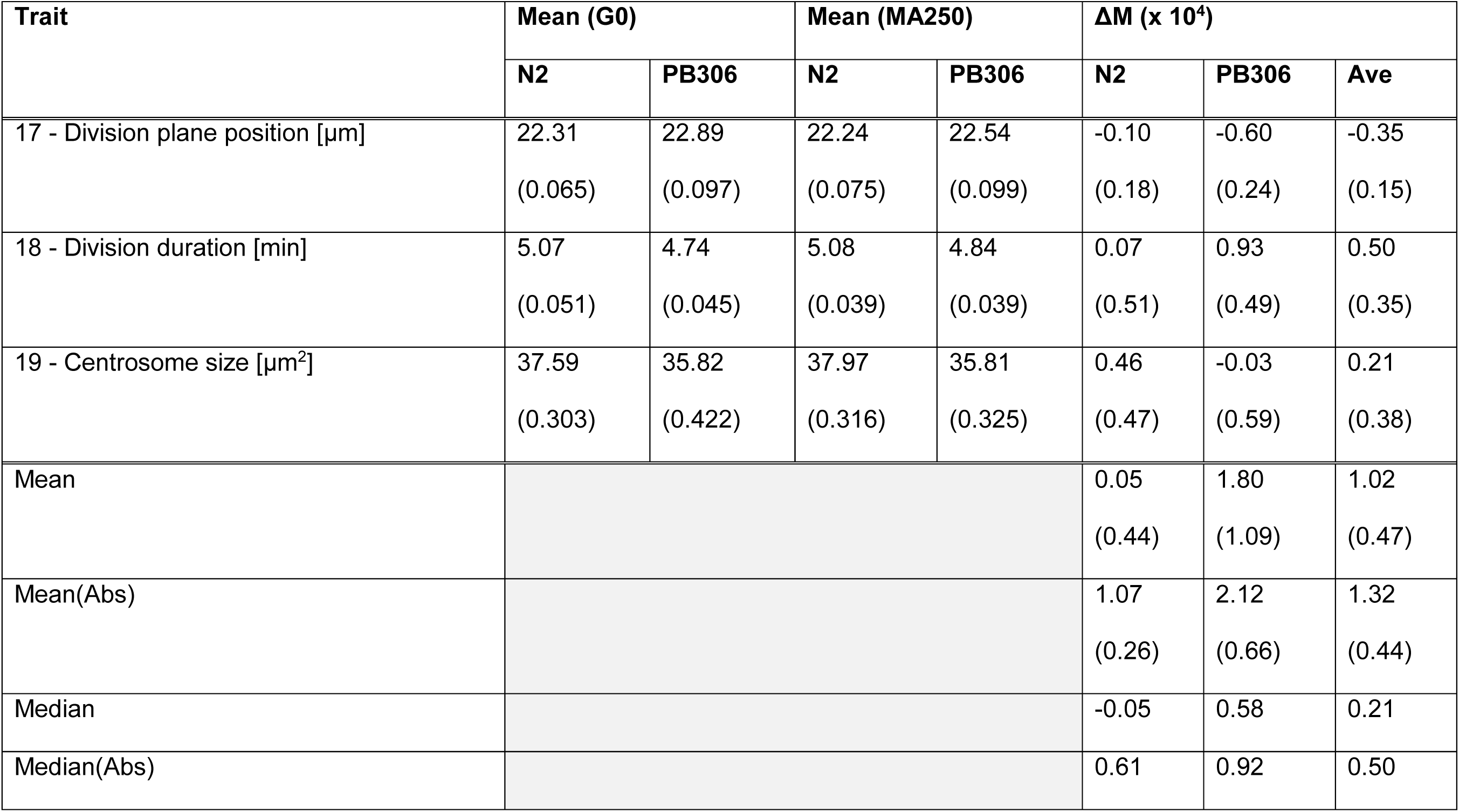
Trait Means, standard errors in parentheses. Column headings are: *Trait*, see Figure 1 for descriptions of traits; *Mean (G0)*, mean trait value of the ancestral G0 control; *Mean (MA250)*, mean of the MA lines; *ΔM*, per-generation change in the trait mean. *ΔM* for traits 9–13 (gray rows) are standardized by the environmental standard deviation rather than the by the mean, thus the per-generation change is given in units of phenotypic standard deviations rather than as a fraction of the mean. Values of *ΔM* marked by * are significantly different from 0 at the experiment-wide P<0.05.

**Figure 1.**
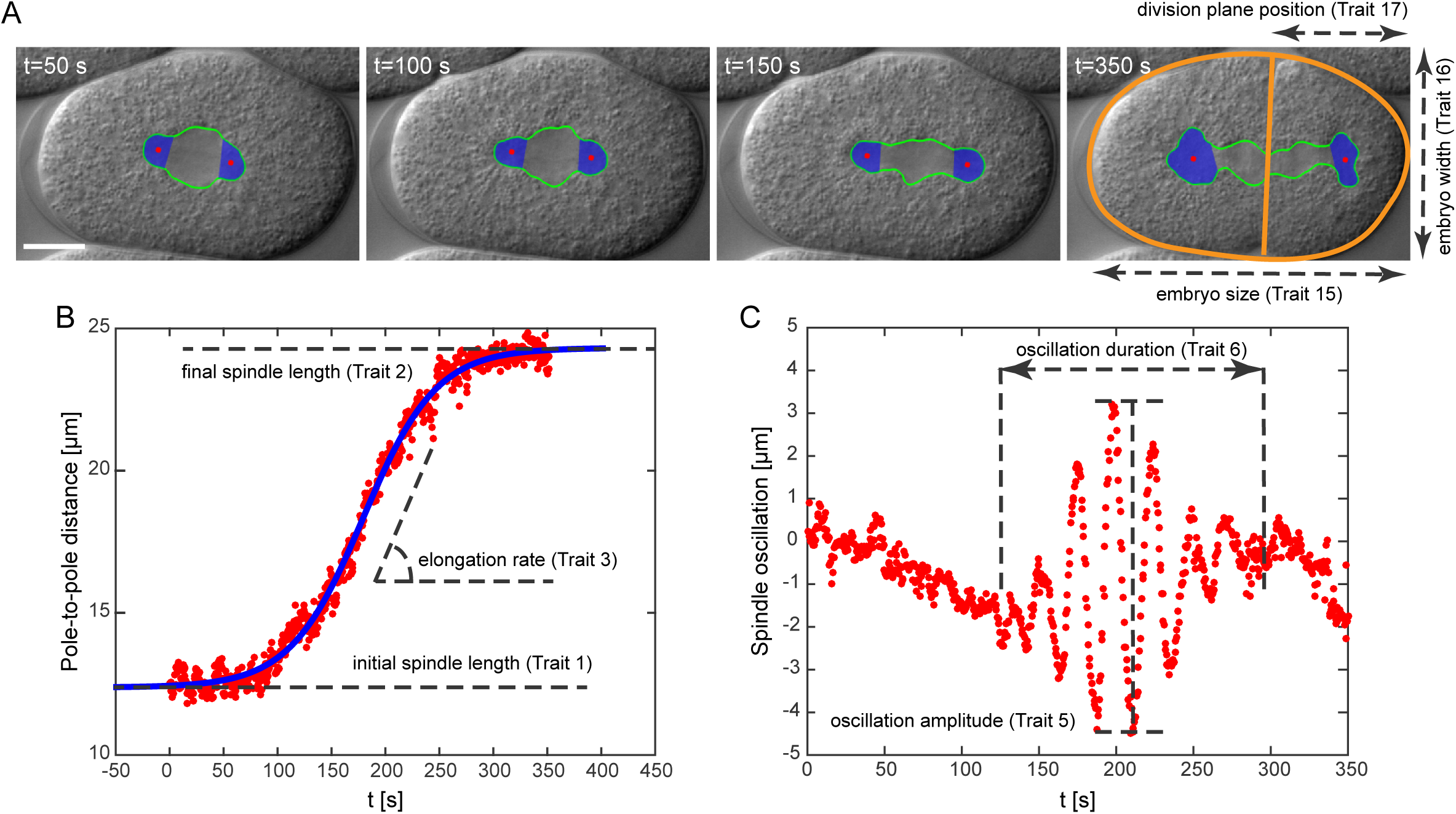
Tracking and measurement of cell-division traits in the first mitotic division of *C. elegans:* (A) Automatic tracking of the spindle (green), centrosomes (blue), cellular boundary (orange, last panel), and position of the division plane (orange, last panel). Measurements for Traits 15–17 are shown in the last panel. Scale bar 10 microns. (B) Pole-to-pole distance as a function of time (red dots). The blue curve is the sigmoid function fitted to the data (see Methods and Materials). Measurements for Traits 1–3 are shown. (C) Spindle oscillation as a function of time (red dots). The distance of the posterior centrosome from the long axis of the embryo is plotted as a function of time. Measurements for Traits 5 and 6 are shown.

The biological particulars of Drosophila and Daphnia are quite different, except that in both species the relevant selection against deleterious mutations is in the heterozygous state. Here we report an analysis of mutational and standing genetic variation in 19 traits related to the first mitotic cell division in the nematode *Caenorhabditis elegans*, chosen on the basis of their relevance to cell biology. The population genetic milieu of *C. elegans* is very different from that of Drosophila or Daphnia. It reproduces predominantly by self-fertilization (Rockman and Kruglyak 2009), so the relevant selection against mutations is in the homozygous state. Moreover, *C. elegans* appears to have undergone one or more global selective sweep(s) within the past 600–1200 generations, resulting in (among other things) linkage disequilibrium that extends over entire chromosomes, little geographic substructure, a large excess of rare alleles (as measured by Achaz’ Y statistic) and a global effective population size on the order of 10^4^ (Andersen *et al*. 2012).

Our primary question of interest is: What is the relationship between VG and VM? Because of the recent history of strong global selection in *C. elegans*, we can imagine two plausible alternative scenarios. First, since genetic variation was recently purged, mutation may predominate: the only genetic variation present is that introduced by new mutations since the recent purge. If so, we should see a strong positive association between VG and VM, and further, VG should consistently be no more than a few hundreds of generations of VM. Alternatively, because LD is so strong and also because the purge of genetic variation was not complete (Thompson *et al*. 2015), idiosyncratic selection at linked loci may predominate, leading to a more or less random association between VG and VM.

Our data permit us to address two additional questions of fundamental importance. First, how consistent are the mutational properties for particular traits in independent trials (i.e., different sets of mutation accumulation lines)? Second, for traits that differ substantially in VM, can we confidently attribute the difference in VM to a difference in the mutational target or a difference in the effects of mutations on the trait? That is, do traits that differ in VM differ because the mutational target is different or because they are differently robust to the effects of mutation (or both)?

## Methods and Materials

*Mutation Accumulation (MA) lines* – The details of the MA lines are reported in Baer *et al*. (2005). Briefly, sets of 100 replicate lines were initiated from highly homozygous populations of the N2 and PB306 strains of *C. elegans* and maintained by serial transfer of a single immature hermaphrodite every generation for 250 generations, at which point each MA line was cryopreserved. The common ancestor (G0) of each set of MA lines was cryopreserved at the outset of the experiment.

*Wild Isolates* - We assayed first cell division in a worldwide collection of 97 wild isolates of *C. elegans*. Strain IDs and collection information are reported in Supplementary Table S1.

*Mitotic Cell Division Phenotype Assays* - The details of maintenance, microscopy, and image processing of the first mitotic spindle in *C. elegans* embryos are reported in Farhadifar and Needleman (2014) and Farhadifar *et al*. (2015). Briefly, all lines were cultured at 24° C on nematode growth media and fed the OP50 strain of *Escherichia coli*. We dissected and imaged the embryos on a 4% agar pad between a slide and coverslip by differential interference contrast (DIC) microscopy. We developed image-processing software to track the spindle poles in the DIC images (Figure 1A; trait numbers are associated with trait descriptions in Table 1). For each embryo, we measured the pole-to-pole distance and fitted a sigmoid function *l(t) = l*_0_ + *l*_1_/(l + exp(−(*t* − *T_l_)/τ))* to the data (Figure 1B). We defined Trait 1, *l*_0_, and Trait 2, *l*_0_ *+ l*_1_, as the initial and final length of the spindle (in *μ*m), respectively (Figure 1B). Trait 3 (the elongation rate of the spindle in *μ*m/minute, see Figure 1B) and Trait 4 (the duration of spindle elongation in seconds) are defined as *l*_1_*/*4*τ* and *τ*, respectively. We quantified spindle oscillation by measuring the distance of the posterior and anterior centrosomes from the long axis of the embryo (Figure 1C). We defined Trait 5 (oscillation amplitude of the posterior centrosome in *μ*m) as the largest peak-to-trough distance of the posterior centrosome (Figure 1C), and Trait 6 (oscillation duration of the posterior centrosome in second) as the total duration of the posterior centrosome oscillates (Figure 1C). We defined Traits 7 and 8 for the anterior centrosome similar to the Traits 5 and 6 for the posterior centrosome. Traits 9 and 11 (in seconds) are defined as the time difference between the mid-spindle elongation (*T_l_*, see above) and the maximum oscillation peak for the posterior and anterior centrosomes, respectively.

Traits 10 and 12 (in seconds) are defined as the time difference between the first oscillation peak and the mid-spindle elongation time for the posterior and anterior centrosomes, respectively. We defined Traits 13 and 14 as the average frequency of centrosome oscillation (in minutes^-1^) of the posterior and anterior centrosomes. Traits 15 and 16 are defined as the length and width of the embryo in *μ*m (Figure 1A last panel). We defined Trait 17 as the position of the division plane from the posterior end of the embryo in *μ*m, and Trait 18 as the duration of the first division in minutes. Trait 19 is defined as the average size of the centrosomes for *t > T_l_ + eτ* (see above).

*Data Analysis* - With 19 traits, there are many more elements in the covariance matrix (190) than there are MA lines (46 or 47), which precludes many standard multivariate quantitative genetic analysis in the absence of some sort of data reduction technique. Because our primary interests relate to the organismal phenotype *per se*, we restrict the analyses to the univariate case, beyond an initial application of multivariate analysis of variance (MANOVA) to address the question of overall variation among lines. As it turns out, the average absolute pairwise correlation between MA line means is not large (mean |*r*| = 0.23), although the range spans nearly the full possible range (range= -0.78, 0.83; Supplementary Table S2). Moreover, the correlations between MA line means are, on average, very similar to the correlations between G0 pseudoline means (mean |*r*| = 0.26, range = -0.66, 0.86).

Data were analyzed for each set of MA lines (N2 and PB306) separately except as noted.

i) *Trait standardization* - Meaningful comparisons among traits require that the traits be standardized on a common scale. Traits can be standardized either by the trait mean, in which case the scaled genetic variance is the squared genetic coefficient of variation, or by the phenotypic standard deviation, in which case the scaled genetic variance is the heritability. The squared genetic CV is naturally related to the evolvability of a trait (Houle 1992; Hansen *et al*. 2011), which is the “opportunity for selection” when the trait is fitness (Crow 1958). In some cases, however, mean standardization is not appropriate, e.g., when the trait value can be either positive or negative. Of the 19 traits, four (Traits 9–12) cannot be meaningfully mean standardized. We report results for raw (unstandardized) data and SD standardized traits from the full data set and results for mean standardized traits from the reduced set of 15 traits. MA lines were mean standardized by dividing each data point by the mean of the G0 ancestor and SD standardized by the square-root of the within-line (environmental) variance averaged over all lines (G0 and MA). Wild isolates were mean standardized by the global mean and SD standardized by the square-root of the within-line variance. In all cases, the important conclusions are qualitatively similar for the two different standardizations.

*ii) Evolution of Trait Means in MA lines (ΔM)* - A directional change in the trait mean over the course of a MA experiment indicates a mutational bias. The per-generation change in the trait mean (*ΔM*) was determined from the slope of the regression of the (standardized) trait mean against generations of MA, i.e., 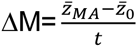 where 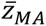 and 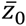 are the means of the MA lines and the G0 ancestors, respectively, and *t* is the number of generations of MA. Regression slopes were calculated from the general linear model

*z_i_=t+Line(Treatment)+Replicate(Line(Treatment))* where *z_i_*, is the standardized value of trait *i, t* is the number of generations of MA (equal to zero for the G0), Line is a random effect representing MA line or G0 pseudoline, Replicate is a random effect representing the experimental unit of observation (i.e., a worm) and Treatment is MA or G0 control. Analyses were performed using the MIXED procedure in SAS v.9.4. Variance components of the random effects were estimated by Restricted Maximum Likelihood (REML) separately for each treatment group using the GROUP= option in the RANDOM and REPEATED statements of the MIXED procedure (Fry 2004). Degrees of freedom were determined by the Kenward-Roger method (Kenward and Roger 1997). Estimation of the regression slope from standardized traits fails to account for sampling variance of the G0 controls, but with the sample sizes in this study (hundreds of control individuals) the bias is negligible, and the empirical 95% confidence intervals calculated by bootstrapping over lines (data not shown) are very close to those calculated from the linear model.

*iii) Mutational Variance (VM)* - The mutational variance is half the difference in the among-line component of variance between the MA lines and the G0 pseudolines, divided by the number of generations of MA, i.e, 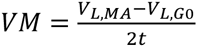, where *V_L,MA_* is the variance among MA lines, *V_L,G_*_0_ is the variance among the G0 pseudolines, and *t* is the number of generations of MA (Lynch and Walsh 1998, p. 330).

As an initial assessment of the overall variation, we considered the multivariate general linear model ***z*** = *Line(Treatment)* + *Replicate(Line(Treatment))*, where ***z*** is the vector of all (standardized) traits and the other variables are as previously defined. Variance components were estimated by REML separately for MA and G0 control groups, and statistical significance of the among-line component of variance was assessed by Likelihood-Ratio Test (LRT) of the model with separate among-line components of variance for MA and G0 groups compared to the model with a single among-line component of variance. The models are nested and differ by a single parameter, so the likelihood ratio is asymptotically chi-square distributed with one degree of freedom.

We repeated the above analysis for each trait individually using the same model and likelihood-ratio test, i.e., *z_i_* = *Gmax* + *Line(Treatment)* + *Replicate(Line(Treatment))*.

*iv) Genetic variance of wild isolates (VG)* - The inferred rate of outcrossing among *C. elegans* in nature is very low (Rockman and Kruglyak 2009), so we treat the wild isolates as if they are homozygous lines. The genetic variance among a set of homozygous lines is half the among-line component of variance (Falconer 1989, p. 265). Variance components were estimated from the linear model *z_i_*, = *Line* + *Replicate(Line)*, where Line represents the wild isolates and z_i_ and Replicate are as above. Significance of the among-line component of variance was assessed by LRT comparison of models with and without the Line term included.

*v) Mutational target (Q_z_) and average squared effect (α^2^*) - The mutational variance, VM, is the product of the genome-wide mutation rate (*U*), the fraction of the genome with the potential to affect the trait (the mutational target, *Q_z_*) and the average squared effect of a mutant allele on the trait, *α^2^*, i.e., VM=*UQ_z_α^2^* (Barton 1990; Kondrashov and Turelli 1992). Changes in trait means (*ΔM*) and variances (VM) and the genomic mutation rate (*U*) can be directly estimated from data, whereas the underlying parameters *Q_z_*, and *α^2^* can only be inferred indirectly. Moreover, in the absence of additional information, *U* and *Q_z_* can only be inferred jointly as the product *UQ_z_* because all combinations of *U* and *Q_z_* that produce the same number of mutations affecting the trait are indistinguishable.

For a selected subset of traits, we assessed the goodness-of-fit of various combinations of the underlying parameters to the data; criteria for our choices of traits and parameter values are explained in the Results. For each combination of parameter values *UQ_z_*, and *α^2^* shown in Supplementary Table S3, we simulated 1000 pseudo-experiments, maintaining the distribution of sample sizes of the actual experiment. The parameters were constrained such that the product 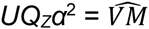, where 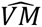 is the observed mutational variance. We investigated two different distributions of mutational effects, the Normal and the Exponential reflected around zero. In the Normal model, mutational effects (*α*) are normally distributed with mean 0 and variance *α^2^*. The simulations from a reflected exponential distribution with variance equal to *α^2^* were obtained by simulating samples from the mixture distribution

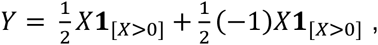
where X is exponentially distributed with rate 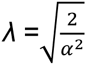 and **1**_[_*_A_*_]_ is the indicator function, which is equal to 1 if *A* and 0 otherwise.

The structure of these models is conceptually related to the House of Cards approximation to the Continuum of Alleles model of mutation (Turelli 1984; Bulmer 1989), in that the effect of an allele after mutation is uncorrelated with the phenotype prior to mutation (although there is no selection in our model). Each simulated MA line is assigned a unique set of mutations that are Poisson distributed among lines with parameter *Ut* where *U* is the haploid genomic mutation rate and *t* is the number of generations of MA. Individual mutations have a non-zero effect on the trait with probability *Q_z_*. Each MA line *i* has a unique genotypic value *g_i_* equal to the sum of its mutational effects drawn from either the Normal or the Reflected Exponential, as described above. Each replicate *j* of MA line *i* is assigned a unique environmental effect *ε_ij_* drawn from a Normal distribution with mean 0 and variance equal to the sum of the within-line variance (VE) and the among-line variance of the G0 controls *(V_L,G0_)*. For each of the eight combinations of the joint parameter *UQ_z_* and *α^2^*, we simulated 1000 samples of line means and compared the distribution of the simulated samples with the observed distribution of line means, using as a metric the empirical estimate of the Kullback-Liebler (KL) divergence (Hausser and Strimmer 2009). The smaller the KL distance, the better the fit. For each of the two DME models we calculated weighted medians of the parameters *UQ_z_* and |*α*|, weighting by the proportion of simulations that a given parameter provided the best fit, given that that model provided the better fit. For example, if the exponential DME provided the better fit in 400 of the 1000 simulations and of those 400, *UQ_z_* = 0.5 was best 200 times, *UQ_z_* = 0.1 was best 150 times and *UQ_z_* = 0.02 was best 50 times and none of the other values of *UQ_z_* was best in any of the 400 replicates, we take the median of 200*0.5+150*0.1+50*0.02 = 0.3 to be the best estimate of *UQ_z_* given that the DME is exponential. We take as the overall best estimate of a parameter to be the weighted mean of the best estimate for each DME model, weighted by the proportion of times that model was better.

*Data Availability* - Data (raw trait values) will be deposited in Dryad upon acceptance of the manuscript. Simulation code (in R) of the DME models is available by request to the authors.

## Results

*i) Evolution of Trait Means in MA lines (ΔΜ)* - Trait means evolved very little over the course of the MA experiment (Table 1). For the N2 lines, the median absolute change in the trait mean for the 15 traits that could be mean-standardized was 0.0034% per-generation; in no case was the change significant at the Bonferroni-corrected experiment-wide 5% level (0.05/(2*15), P<0.0017). For the PB306 lines, the median absolute change was 0.0073% per-generation, and only two traits (1 and 4) changed significantly at the experiment-wide 5% level. *ΔΜ* was not significantly correlated between the two sets of MA lines (*r* = 0.15, P>0.60). Of the 19 traits, nine changed in the same direction in both sets of lines and ten changed in opposite directions, exactly as predicted if the direction of change was random. Moreover, these results are consistent with the traits being under some degree of stabilizing selection (perhaps collectively; Farhadifar *et al*. 2015), because deleterious mutations do not have consistently directional effects. The *ΔM*s for these traits can be compared to *ΔΜ* for other traits expected to be under directional selection. For example, in these same sets of lines lifetime reproduction weighted by probability of survival (“Total fitness”) decreased by about 0.1% per-generation (Baer *et al*. 2006) and body volume at maturity decreased by about 0.07% per generation (Ostrow *et al*. 2007); in each case the change was highly significant and consistent between the two sets of lines.

*ii) Mutational Variance (VM)* - Integrated over all 19 SD-standardized traits, MANOVA revealed highly significant accumulation of mutational variance in both sets of MA lines (N2, LRT chisquare = 35.3, df = 1, P<<0.0001; PB306, LRT chi-square = 28.4, df = 1, P<<0.0001). However, in both sets of lines MANOVA also revealed a highly significant among-line component of variance in the ancestral G0 controls, and trait-by-trait analysis revealed many cases in which the among-line variance of the G0 controls was marginally significant (Table S4). There are several potential, not mutually-exclusive explanations for non-zero among-line variance of the G0, including residual genetic variation, heritable cross-generational environmental effects (Baugh 2013; Jobson *et al*. 2015), and line-by-environment correlations in the experimental protocol. The sampling design of these experiments constitutes a kind of “anti-Goldilocks zone” in the context of the likelihood-ratio test, whereby there are sufficiently many G0 pseudolines for the REML estimates of the variance component to be non-zero but too few pseudolines to provide much power for the LRT of the model with the among-line component of variance estimated separately for the two treatments against the model with a single among-line component of variance. For none of the 38 (2×19) strain/trait combinations was the among-line variance of the G0 controls significant at the Bonferroni-corrected level, and in only one case (trait 18 in the N2 lines) was the uncorrected P < 0.01. In contrast, the among-line component of variance of the MA lines was significant at the experiment-wide 5% level in almost all cases, failing to reach significance only for trait 13 in the PB306 lines and trait 14 in both sets of lines. More convincingly, in 37/38 cases the point estimate of the among-line variance of the MA lines was greater than that of the G0 pseudolines, the sole exception being trait 13 in the PB306 lines. Thus, we proceed under the assumption that the point estimates of VM represent a reasonable approximation of the truth, even though they do not reach experiment-wide 5% significance in most cases.

Averaged over the two sets of MA lines, mean-standardized VM varies by slightly under two orders of magnitude, from 3.3 × 10^−7^/generation for trait 14 to 2.6 × 10^−5^/generation for trait 7 (Table 2). These values can be put into context by comparison to a set of life history traits measured in these same sets of MA lines (Supplementary Table S5). The average VM for the spindle traits (mean = 6 × 10^−6^/gen, median = 2 × 10^−6^/gen) is substantially smaller than that for the life history traits (mean = 9 × 10^−5^/gen, median = 8 × 10^−5^/gen), although the ranges of variability overlap.

**Table 2.** Variances of G0 mean-standardized traits (= squared coefficient of variation). Standard errors in parentheses. Column headings are: *VL,G0*, among-line variance of G0 pseudolines; *VL,MA*, among-line variance of MA lines; *VE,G0*, within-line variance of G0 pseudolines; *VE,MA*, within-line variance of MA lines; *VM*, mutational variance (× 10^6^); *ave VM*, average VM of the two strains. Standard errors of VM for individual traits are calculated from the square-root of the sum of the sampling variances of the G0 pseudolines and MA lines. Standard errors of the among-trait mean VM are calculated as the among-trait variance divided by the square-root of the number of traits.

| **Trait** | **Strain** | **VL,G0 (x 10<sup>4</sup>)** | **VL,MA (x 10<sup>4</sup>)** | **VE,G0 (x 10<sup>4</sup>)** | **VE,MA (x 10<sup>4</sup>)** | **VM (x 10<sup>6</sup>)** | **ave VM (x 10<sup>6</sup>)** |
| --- | --- | --- | --- | --- | --- | --- | --- |
| 1 | N2 | 2.52 (1.44) | 3.30 (1.13) | 48.52 (3.15) | 47.03 (2.13) | 0.16 (0.37) | 0.42 (0.62) |
|  | PB | 9.73 (4.73) | 13.16 (3.51) | 60.28 (4.50) | 72.59 (3.29) | 0.69 (1.18) |  |
| 2 | N2 | 2.47 (1.18) | 3.60 (1.02) | 24.24 (1.57) | 23.26 (1.06) | 0.23 (0.31) | 0.64 (0.28) |
|  | PB | 2.28 (1.25) | 7.56 (1.99) | 23.31 (1.74) | 35.47 (1.61) | 1.06 (0.47) |  |
| 3 | N2 | 9.48 (6.20) | 50.45 (12.89) | 229.00 (14.88) | 234.90 (10.66) | 8.19 (2.86) | 10.18 (2.13) |
|  | PB | 0 | 60.86 (15.85) | 209.80 (15.34) | 260.50 (11.85) | 12.17 (3.17) |  |
| 4 | N2 | 30.67 (15.01) | 47.83 (13.77) | 346.30 (22.50) | 372.90 (16.93) | 3.43 (4.07) | 5.26 (2.55) |
|  | PB | 5.50 (7.56) | 40.98 (13.35) | 327.40 (24.43) | 372.20 (16.95) | 7.10 (3.07) |  |
| 5 | N2 | 5.63 (5.57) | 52.44 (14.28) | 238.90 (18.75) | 299.40 (13.94) | 9.36 (3.07) | 8.99 (1.95) |
|  | PB | 0 | 43.04 (12.06) | 265.00 (20.17) | 308.40 (13.98) | 8.61 (2.41) |  |
| 6 | N2 | 22.83 (13.42) | 36.39 (12.18) | 405.90 (26.81) | 410.20 (19.11) | 2.71 (3.62) | 5.96 (2.85) |
|  | PB | 11.11 (12.34) | 57.13 (18.22) | 451.70 (35.19) | 437.90 (19.95) | 9.20 (4.40) |  |
| 7 | N2 | 60.82 (31.62) | 207.70 (52.44) | 825.50 (54.43) | 822.60 (38.26) | 29.38 (12.25) | 25.97 (8.79) |
|  | PB | 68.62 (40.18) | 181.40 (48.66) | 875.20 (68.00) | 914.60 (41.52) | 22.56 (12.62) |  |
| 8 | N2 | 42.13 (25.68) | 130.10 (37.58) | 850.40 (56.08) | 939.00 (43.69) | 17.59 (9.10) | 22.99 (7.60) |
|  | PB | 41.14 (30.68) | 183.10 (52.51) | 964.40 (74.87) | 1115.0 (50.63) | 28.39 (12.16) |  |
| 13 | N2 | 5.48 (2.56) | 11.28 (2.97) | 49.77 (3.29) | 54.92 (2.56) | 1.16 (0.78) | 0.58 (0.60) |
|  | PB | 8.83 (4.34) | 3.82 (1.43) | 57.95 (4.51) | 57.76 (2.62) | 0.00 (0.91) |  |
| 14 | N2 | 3.47 (2.05) | 6.25 (2.15) | 61.49 (4.06) | 74.01 (3.46) | 0.56 (0.59) | 0.33 (0.41) |
|  | PB | 2.93 (2.36) | 3.46 (1.52) | 70.58 (5.50) | 73.93 (3.36) | 0.11 (0.56) |  |
| 15 | N2 | 1.59 (0.80) | 3.27 (0.90) | 20.25 (1.29) | 21.30 (0.95) | 0.34 (0.24) | 0.81 (0.25) |
|  | PB | 1.62 (0.88) | 8.07 (1.97) | 17.85 (1.31) | 30.63 (1.33) | 1.29 (0.43) |  |
| 16 | N2 | 2.37 (1.41) | 4.50 (1.39) | 50.36 (3.22) | 44.81 (2.00) | 0.43 (0.40) | 0.59 (0.33) |
|  | PB | 2.59 (1.85) | 6.35 (1.82) | 61.88 (4.55) | 51.78 (2.24) | 0.75 (0.52) |  |
| 17 | N2 | 0.31 (0.60) | 3.38 (1.12) | 39.57 (2.53) | 134.70 (5.99) | 0.61 (0.25) | 0.81 (0.24) |
|  | PB | 1.44 (1.05) | 6.51 (1.80) | 32.49 (2.39) | 46.32 (2.00) | 1.01 (0.42) |  |
| 18 | N2 | 15.63 (6.40) | 21.57 (5.81) | 82.81 (5.28) | 134.70 (5.99) | 1.19 (1.73) | 2.28 (1.18) |
|  | PB | 7.77 (4.74) | 24.64 (6.42) | 118.10 (8.61) | 131.10 (5.73) | 3.37 (1.60) |  |
| 19 | N2 | 5.38 (4.23) | 24.84 (7.05) | 191.80 (12.13) | 179.20 (7.92) | 3.89 (1.64) | 3.54 (1.42) |
|  | PB | 11.23 (8.04) | 27.20 (8.25) | 255.60 (18.39) | 253.70 (10.91) | 3.19 (2.30) |  |
| Mean | N2 |  |  |  |  | 5.28 (2.20) | 5.96 (2.19) |
|  | PB |  |  |  |  | 6.60 (2.30) |  |
| Median | N2 |  |  |  |  | 1.19 | 2.28 |
|  | PB |  |  |  |  | 3.19 |  |

In contrast to the *ΔMs*, which are uncorrelated between the two sets of MA lines, the mutational variances are highly correlated between the N2 and the PB306 lines. For the full data set of 19 traits, the correlation between the raw (unstandardized) VMs in the two strains is 0.95 (P<0.00001; Figure 2) and the correlation for the subset of 15 mean-standardized traits is 0.89 (P<0.0001; Supplementary Figure S1). The correlation between the mutational heritabilities in the two strains is smaller, although still significantly positive (*r* = 0.56, P<0.02).

**Figure 2.**
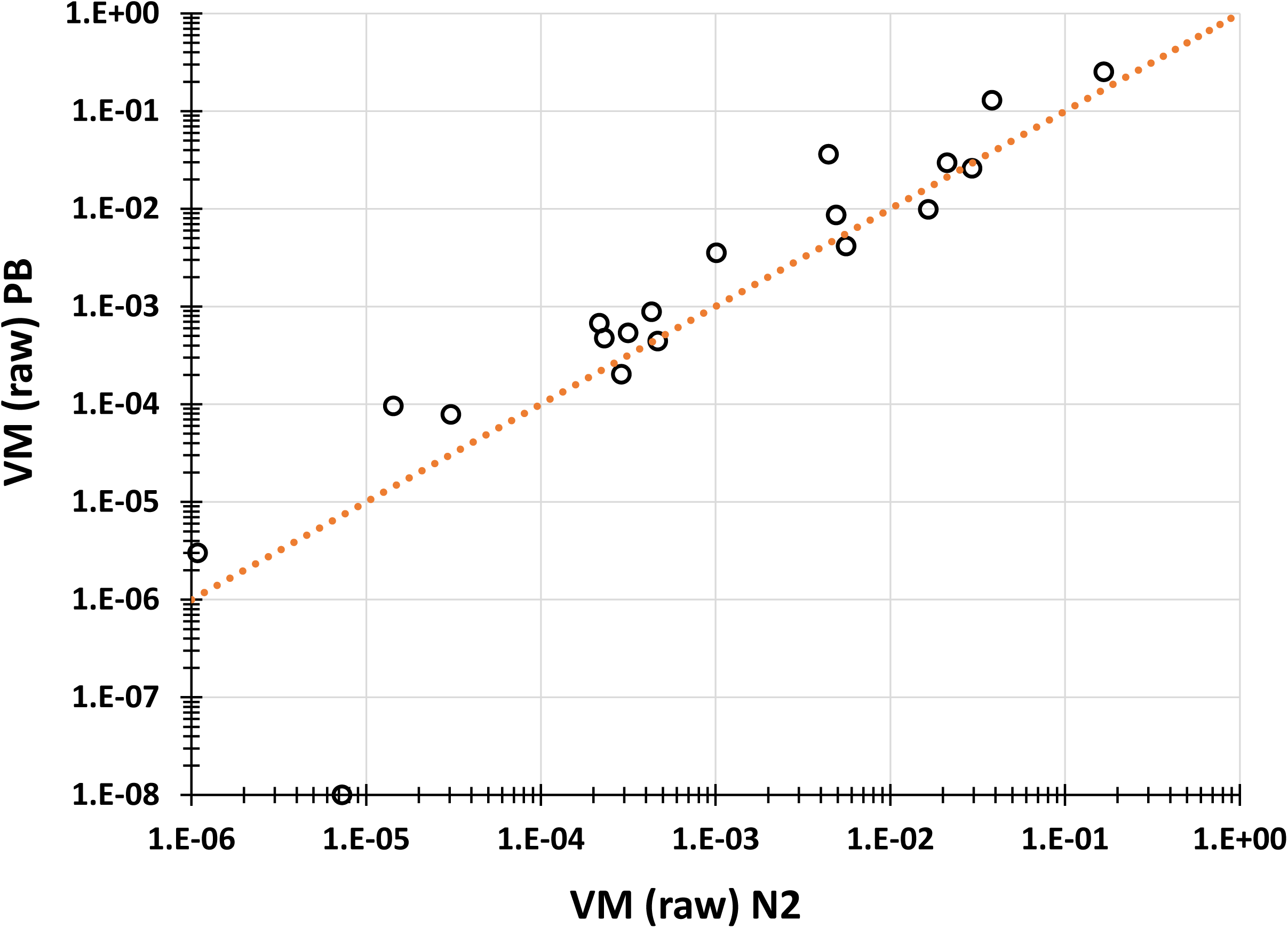
Raw VM (PB306) plotted against raw VM (N2). The dashed line represents the line of equality.

Averaged over both sets of MA lines, there is a strong positive correlation between VM and VE (*r* ≈ 0.95). This result is commonly observed, and is consistent with the idea that genetic variation and environmental variation have a common biochemical and/or physiological basis (Meiklejohn and Hartl 2002).

*iii) Genetic variance of wild isolates (VG)* - For all traits the among-line component of variance (raw and mean-standardized) among the wild isolates is highly significantly different from zero (P<0.0001 in all cases), as is the broad-sense heritability, *H^2^* (Table 3). However, the potentially non-zero among-line variance of the G0 ancestors of the MA lines introduces the possibility that some fraction of the among-line variance of the wild isolates is not true genetic variance. To address that possibility, we subtracted the average of the two estimates of the among-line variance of the G0 controls from the estimate of the among-line variance of the wild isolates before calculating VG; we refer to the corrected estimate of VG as VG^*^ (Table 3). On average, VG^*^ is reduced by about 20–30% relative to the uncorrected VG (mean reduction = 27%; median reduction = 20%). For the full set of 19 unstandardized traits, the correlation between the average mutational variance VM and the genetic variance VG^*^ is nearly perfect (*r* = 0.99, P<0.0001; Supplementary Figure S2); the correlation is essentially the same for the 15 mean-standardized traits (*r* = 0.95, P<0.0001; Figure 3). The correlation between the mutational heritability and *H^2^* is somewhat smaller but remains highly significant (*r* = 0.69, P<0.002).

**Table 3.** Summary statistics of wild isolates, standard errors in parentheses. Column headings are: *Mean*, trait mean; *VG (raw)*, standing genetic variation calculated from the raw data; *VE (raw)*, environmental (within-strain) variance calculated from the raw data; *H^2^*, broad-sense heritability; VG^*^, mean-standardized VG corrected by subtracting the average among-line variance of the MA controls; *t_P_ (raw)*, expected persistence time of a new mutation (*t_P_*=VG/VM) calculated from raw data corrected by subtracting average among-line variance of MA controls from VG; *t_P_ (std)* calculated from mean-standardized data.

| *Trait* | *Mean* | *VG (raw)* | *VE (raw)* | *H<sup>2</sup>* | *VG\* (std)*<br>*(x 10<sup>3</sup>)* | *t<sub>p</sub>\* (raw)* | *t<sub>p</sub>\* (std)* |
| --- | --- | --- | --- | --- | --- | --- | --- |
| 1 | 11.61 (0.04) | 0.06 (0.01) | 0.74 (0.02) | 0.07 (0.01) | 0.17 (0.24) | 419 (641) | 394 (611) |
| 2 | 24.89 (0.08) | 0.25 (0.04) | 1.40 (0.04) | 0.13 (0.02) | 0.28 (0.13) | 414 (189) | 441 (219) |
| 3 | 5.76 (0.07) | 0.20 (0.03) | 0.80 (0.02) | 0.17 (0.03) | 5.71 (1.24) | 544 (150) | 561 (149) |
| 4 | 35.67 (0.38) | 6.24 (1.03) | 43.56 (1.29) | 0.11 (0.02) | 3.91 (1.36) | 763 (386) | 743 (390) |
| 5 | 7.38 (0.09) | 0.40 (0.06) | 1.45 (0.04) | 0.18 (0.03) | 7.16 (1.41) | 878 (231) | 797 (214) |
| 6 | 126.4 (1.4) | 78.50 (13.27) | 629.0 (18.7) | 0.10 (0.02) | 4.01 (1.47) | 781 (432) | 674 (351) |
| 7 | 3.49 (0.08) | 0.26 (0.04) | 1.03 (0.03) | 0.17 (0.03) | 18.07 (5.14) | 953 (354) | 696 (269) |
| 8 | 105.6 (1.7) | 128.8 (21.4) | 878.0 (26.1) | 0.11 (0.02) | 9.54 (3.33) | 524 (201) | 415 (161) |
| 9 | -16.89 (0.60) | 13.87 (2.50) | 157.6 (4.8) | 0.07 (0.01) |  | 864 (529) |  |
| 10 | 34.30 (0.77) | 21.70 (4.15) | 317.7 (9.7) | 0.06 (0.01) |  | 703 (359) |  |
| 11 | -32.65 (0.62) | 11.45 (2.68) | 330.8 (10.1) | 0.03 (0.01) |  | 448 (210) |  |
| 12 | 15.41 (0.71) | 16.59 (3.54) | 343.0 (10.5) | 0.04 (0.01) |  | 599 (517) |  |
| 13 | 2.75 (0.01) | 0.004 (0.001) | 0.037 (0.001) | 0.09 (0.02) | 0.19 (0.27) | 404 (533) | 335 (384) |
| 14 | 2.62 (0.01) | 0.003 (0.001) | 0.046 (0.001) | 0.05 (0.01) | 0.26 (0.19) | 708 (1147) | 778 (989) |
| 15 | 50.96 (0.12) | 0.56 (0.10) | 5.17 (0.15) | 0.09 (0.02) | 0.13 (0.08) | 165 (65) | 163 (66) |
| 16 | 34.00 (0.10) | 0.35 (0.07) | 5.81 (0.17) | 0.05 (0.01) | 0.18 (0.14) | 322 (211) | 304 (195) |
| 17 | 22.38 (0.06) | 0.16 (0.03) | 2.10 (0.06) | 0.07 (0.01) | 0.27 (0.10) | 325 (116) | 330 (121) |
| 18 | 4.83 (0.03) | 0.04 (0.01) | 0.30 (0.01) | 0.12 (0.02) | 1.31 (0.59) | 537 (320) | 575 (326) |
| 19 | 37.17 (0.38) | 6.33 (1.00) | 31.08 (0.88) | 0.14 (0.02) | 4.02 (1.00) | 1194 (505) | 1135 (494) |
| Mean |  |  |  | 0.10 (0.01) | 3.68 (1.29) | 608 (59) | 556 (65) |
| Median |  |  |  | 0.09 | 2.61 | 544 | 561 |

**Figure 3.**
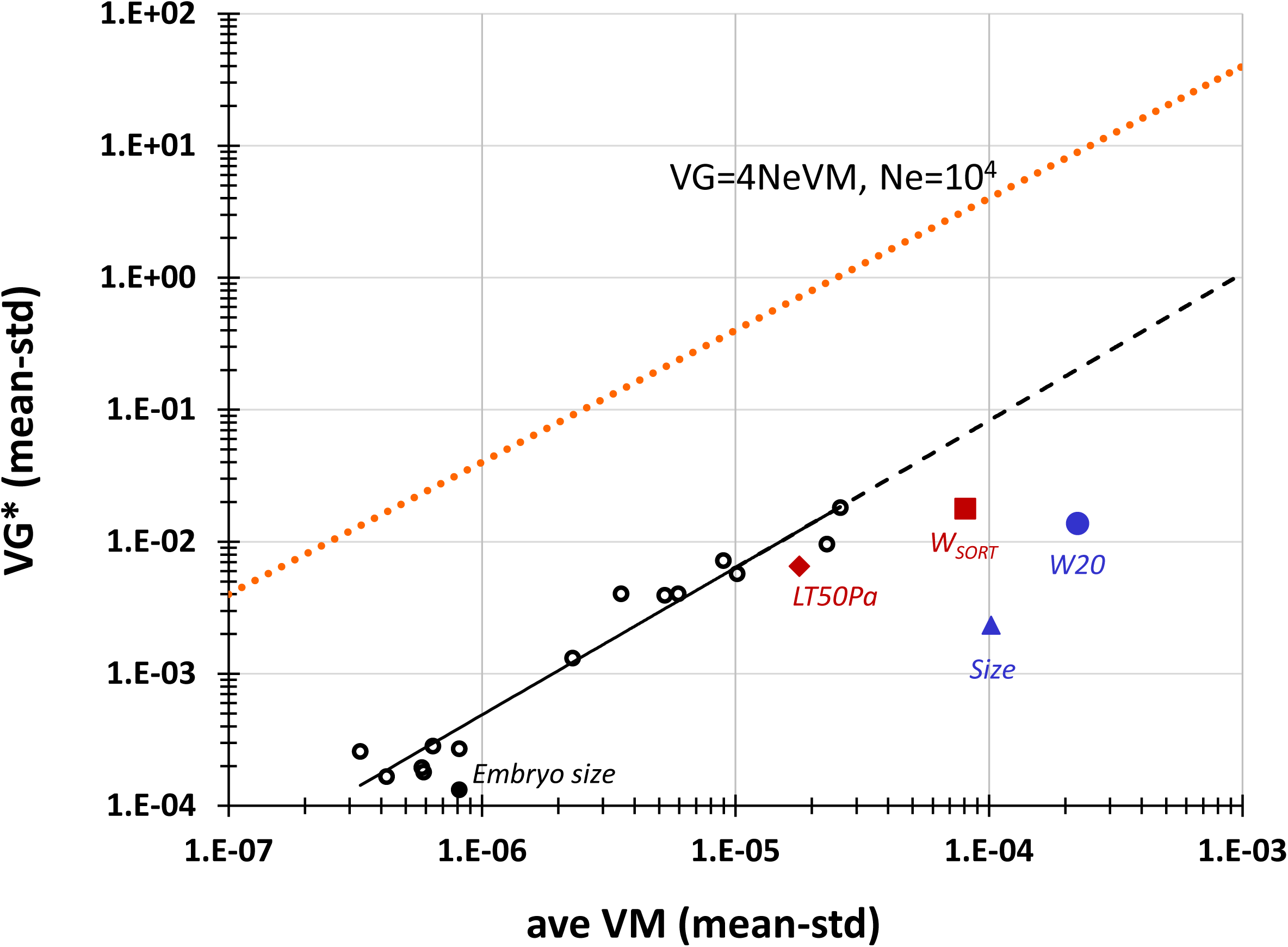
Mean-standardized VG^*^ plotted against mean-standardized VM. The solid black line shows the best-fit of the spindle trait data; the dashed black line represents the extension of the best-fit line. The orange dashed line shows 4*N_e_*VM for *N_e_* = 10^4^. See text for description of labeled traits and experimental details. Traits labeled in orange were measured on the same set of wild isolates included in this study.

The ratio VG/VM has several interpretations, depending on the context. First, in an infinite population at MSB, it represents the persistence time *(t_P_)* of a new mutation, i.e., the expected number of generations before the mutant allele is removed by selection (Garcia-Dorado *et al*. 2003). The stronger selection is, the shorter the persistence time. Second, for a neutral trait in a finite population, VG = *2NeVM* at mutation-drift equilibrium (Lynch and Hill 1986), so VG/VM is equal to *2Ne (4Ne* in the case of obligate self-fertilization, which is approximately the case with *C. elegans)*. Finally, VG/VM represents the number of generations of mutation required to produce a given amount of genetic variance, irrespective of other evolutionary forces.

For almost all of the traits in this study, the ratio VG^*^/VM (called *t_P_^*^* in Table 3) falls within the relatively narrow window of 300–800. Two traits are obvious outliers: Embryo size (Trait 15; *t_P_^*^ ≈* 160) and Centrosome size (Trait 19; *t_P_^*^* > 1100). Embryo size has been previously inferred to be under long-term stabilizing selection (Farhadifar *et al*. 2015) and the reduced *t_P_* is consistent with stronger selection on that trait than on the other traits. We have no intuition about why Centrosome size is a high outlier. Balancing selection for some unknown reason is possible, although random chance seems as plausible as anything.

*iv) Mutational target (Q_z_) and average-squared effect (α^2^)* - Ultimately, we would like to understand the underlying genetic causes of differences in VM between traits - if VM differs substantially between two traits, is it because different numbers (or kinds) of loci contribute to variation, or is it because the average effects of mutations are different? In other words, is the difference due to different mutational targets, or different mutational robustness? As a first pass at this question, we chose two traits, Anterior centrosome oscillation duration (trait 8) and Division plane position (trait17), that met the following criteria: they are among the most highly and consistently divergent in mean-standardized VM (~ 28X in both strains), the G0 means do not differ significantly between the two strains, the trait means did not evolve significantly over the course of the experiment *(ΔM ≈* 0), and the distribution of MA line means is approximately symmetrically distributed around zero. These properties allow us to pool the data from the two sets of MA lines. For these purposes, the resulting data set of 93 250-generation MA lines, replicated on average ~25X, provides a uniquely large, well-replicated data set in a multicellular eukaryote.

The parameters of our distribution of mutational effects (DME) model are the joint parameter *UQ_z_*, where *U* is the genome-wide mutation rate and *Q_z_* is the fraction of the genome that potentially affects the trait, and the average squared mutational effect (*α^2^*), subject to the constraint 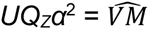. Data on *U* in *C. elegans* that include all classes of mutations (single nucleotide variants, indels, structural variants, copy number variants, TEs) remain incomplete, but there are good reasons to believe *U* is not less than about 0.5 and probably not more than about two per haploid genome per generation (Denver *et al*. 2004; Phillips *et al*. 2009; Lipinski *et al*. 2011; Denver *et al*. 2012). We allowed the joint parameter *UQ_z_* to vary over six orders of magnitude, from 0.5 down to 6.4 × 10^−6^, resulting in a mutational target from half the genome down to 0.00064% of the genome. For each value of *UQ_z_, α^2^* was then chosen so as to give the observed VM.

The simulation parameters are given in Supplementary Table S3, and the results are summarized in Table 4 and Supplementary Table S3. The first obvious conclusion is that even for as large a data set as this one, combinations of mutational parameters *(UQ_z_* and *α^2^*) spanning several orders of magnitude can produce results consistent with the data (Supplementary Table S3), which in turn means that the same set of mutational parameters can lead to very different VM. The likely explanation for this evident lack of power is that although traits differ considerably in mean-standardized VM (range ~80X over the 15 mean-standardized traits, 28X for this pair of traits), they differ by much less in mutational heritability (range < 10X over all 19 traits, approximately equal for this pair of traits). Noise introduced by VE overpowers the signal of VM and, worse, the noise is highly correlated with the signal.

**Table 4.**
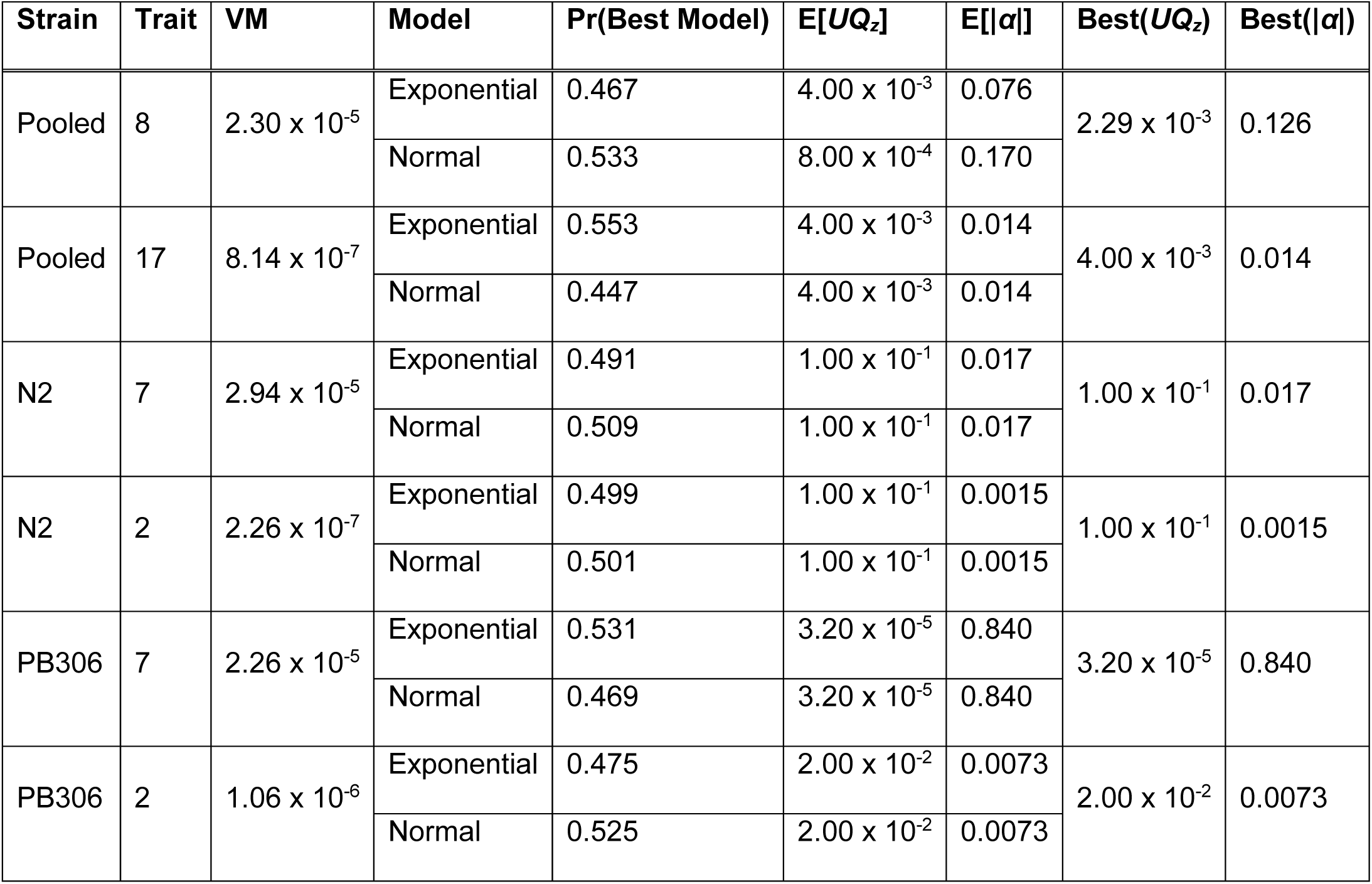
Summary of Goodness-of-Fit tests of mutation models (see text for details). Column headings are: Strain (N2, PB306, or pooled); Trait (see Figure 1 and text for definitions); VM, mean-standardized mutational variance; Model (Exponential or Normal); Pr(Best Model), the proportion of simulation replicates in which the model provided the better fit; E[*UQ_z_*], median target size weighted by the proportion of simulation replicates in which the target provided the best fit given that the model provided the better fit; E[|*α*|], median square-root of the average squared effect, weighted by the proportion of replicates in which the average effect provided the best fit given that the model provided the better fit; Best(*UQ_z_*), mean target weighted by the proportion of replicates that the model provided the better fit; Best(|*α*|), mean average absolute effect, weighted by the proportion of replicates that the model provided the better fit.

| Strain | Trait | VM | Model | Pr(Best Model) | $E[UQ_z]$ | $E[\vert \alpha\vert ]$ | Best($UQ_z$) | Best($\vert \alpha\vert $) |
| --- | --- | --- | --- | --- | --- | --- | --- | --- |
| Pooled | 8 | $2.30 \times 10^{-5}$ | Exponential | 0.467 | $4.00 \times 10^{-3}$ | 0.076 | $2.29 \times 10^{-3}$ | 0.126 |
| | | | Normal | 0.533 | $8.00 \times 10^{-4}$ | 0.170 | | |
| Pooled | 17 | $8.14 \times 10^{-7}$ | Exponential | 0.553 | $4.00 \times 10^{-3}$ | 0.014 | $4.00 \times 10^{-3}$ | 0.014 |
| | | | Normal | 0.447 | $4.00 \times 10^{-3}$ | 0.014 | | |
| N2 | 7 | $2.94 \times 10^{-5}$ | Exponential | 0.491 | $1.00 \times 10^{-1}$ | 0.017 | $1.00 \times 10^{-1}$ | 0.017 |
| | | | Normal | 0.509 | $1.00 \times 10^{-1}$ | 0.017 | | |
| N2 | 2 | $2.26 \times 10^{-7}$ | Exponential | 0.499 | $1.00 \times 10^{-1}$ | 0.0015 | $1.00 \times 10^{-1}$ | 0.0015 |
| | | | Normal | 0.501 | $1.00 \times 10^{-1}$ | 0.0015 | | |
| PB306 | 7 | $2.26 \times 10^{-5}$ | Exponential | 0.531 | $3.20 \times 10^{-5}$ | 0.840 | $3.20 \times 10^{-5}$ | 0.840 |
| | | | Normal | 0.469 | $3.20 \times 10^{-5}$ | 0.840 | | |
| PB306 | 2 | $1.06 \times 10^{-6}$ | Exponential | 0.475 | $2.00 \times 10^{-2}$ | 0.0073 | $2.00 \times 10^{-2}$ | 0.0073 |
| | | | Normal | 0.525 | $2.00 \times 10^{-2}$ | 0.0073 | | |

Nevertheless, some signal does emerge. Comparison of trait 8 (high VM, ≈ 2×10^−5^/generation) and trait 17 (low VM, ≈ 8×10^−7^/generation) shows that the best-fit mutational targets are not very different between the low VM trait 17 (Best(*UQ_z_*) = 0.050) and the high VM trait 8 (Best(*UQ_z_*) = 0.058), whereas the best fit absolute average effect is approximately five-fold greater for trait 8 than for trait 17 (Best(|*α*|) = 0.109 v. 0.020). Neither the Reflected Exponential nor the Normal model was consistently better.

A second pair of traits, the high-VM trait 7 (Anterior centrosome oscillation amplitude) and the low-VM trait 2 (Spindle final length), also met our criteria except that the G0 means differ significantly between N2 and PB306. We therefore repeated the preceding analysis separately for each of the two sets of MA lines. These comparisons reinforce the inference that differences in effect size have more influence on differences in VM than do differences in target size. In both sets of lines, best-fit mutational targets are either similar or *smaller* for the high VM trait 7 than for the low VM trait 2, whereas best-fit average effects are larger for the high VM trait 7, much larger in the PB306 lines (Table 4).

## Discussion

Two robust conclusions emerge from this study, which have considerable significance in the larger context of evolutionary genetics. First, for this relatively large set of functionally-related but (on average) only modestly correlated traits, the mutational process is highly repeatable: the correlation between estimates of trait-specific VM in two independent sets of MA lines derived from different ancestors is ~ 0.9.

The high repeatability of the mutational process was hardly a foregone conclusion. To put this result in perspective, consider the contrast with fitness-related traits in *Drosophila melanogaster*, which are so noisy and inconsistent that an influential Perspectives piece in *Genetics* in 1999 was subtitled the “Riddle of Deleterious Mutation” (Keightley and Eyre-Walker 1999). The “riddle” being why estimates of mutational parameters for fitness-related traits in *D. melanogaster* are so noisy and inconsistent. There are probably several factors at play, including the demonstrable genetic variation for mutation rate in *D. melanogaster* (Schrider *et al*. 2013). Although we have yet to exhaustively characterize the mutational process in these two strains of *C. elegans* for all categories of molecular mutations, the base-substitution (Denver *et al*. 2012; F. Besnard and M-A. Felix, personal communication) and short-tandem repeat (Phillips *et al*. 2009) mutation rates are quite similar in the two strains.

We believe that one key factor underlying the consistency of the results of this study is also the simplest: experimental consistency. The mutation accumulation lines were maintained in the same lab at the same time under the same conditions, and the phenotypic assays were done in the same lab by the same person at the same time under the same conditions. Further, the level of replication in these experiments is substantially greater (~25 replicates per line) than in many, albeit not all, phenotypic assays of MA lines. This is especially important because the mutational heritabilities for these traits are actually quite low (VM/VE ≈10^−4^; Supplementary Table S4; compare to values in Table 1 of Houle *et al*. (1996)).

It is certainly possible that life-history traits are somehow qualitatively different than the traits in this study. Our traits are restricted to a single, narrow window of time in development, so the phenotype, and thus the phenotypic variance, is not integrated over a long period. We have previously assayed lifetime productivity and size at maturity in these same lines. Averaged over six assays at two temperatures, VM for G0 mean-standardized lifetime productivity varies by less than threefold between the two strains (data from Table 2 of Baer *et al*. (2006)); size at maturity varies by 1.5-fold (data from Table 2 of Ostrow *et al*. (2007)). Those values are well within the range of variation between the two strains for single traits in this study. In contrast, VM for egg-to-adult viability in *D. melanogaster* varies by at least 27-fold across studies, and VM for abdominal bristle number varies by at least 130-fold (data from Table 1 of Houle *et al*. (1996)). Thus, the difference in repeatability between this study and the Drosophila oeuvre does not seem to be due to a qualitative difference between categories of traits.

The second robust result is that VM almost perfectly predicts VG for these traits (Figure 3, Supplementary Figure S2). Again, this was not a foregone conclusion (Charlesworth 2015). This finding is not without precedent, however, as evidenced by Figure 1 of Houle (1998). Houle calculated a correlation between VG and VM of 0.95 for eight life-history and morphological traits in *D. melanogaster*. Lynch *et al*. (1998) reported similar data for nine life-history and morphological traits in *Daphnia pulex*, although they did not explicitly calculate the correlation between VG and VM (*r* = 0.75, reanalysis of data in their Tables 1 and 3). An analogous relationship between VM and between-species divergence was reported for a set of several thousand gene-expression traits in *D. melanogaster* (Rifkin *et al*. 2005); the correlation between VM and between-species divergence ranged between 0.25 and 0.4 (P<0.0001) for three species pairs. Similar data exist for other sets of traits and in other organisms, and we predict the correlation between mean-standardized VM and VG will generally be large and positive.

There are two, potentially interrelated underlying evolutionary mechanisms that predict a large positive correlation between VG and VM. The first is the interplay between mutation and random genetic drift. For a neutral trait at mutation-drift equilibrium (MDE) in a selfing organism, VG = *4N_e_VM* (Lynch and Hill 1986). Global *N_e_* of *C. elegans* has been estimated from the standing nucleotide polymorphism (*θ*) to be on the order of 10^4^ (Andersen *et al*. 2012). In no case does VG of any of the traits investigated here come close to the value of 40,000VM predicted for a neutral trait at MDE (the dashed line in Figure 3); the average is about 550VM. However, *C. elegans* is almost certainly far from global MDE, so it seems intuitively obvious that VG should be well below the value predicted at MDE, even for a neutral trait. However, both VG and *θ* increase at a rate proportional to the mutation rate and decrease by drift at a rate inversely proportional to *N_e_*. It is definitely possible that the loci that underlie most quantitative genetic variation mutate 70-fold more slowly than do single nucleotides. It seems less likely that *N_e_* differs consistently by that much between the two categories of loci.

More importantly, it seems very unlikely to us that these traits are neutral over the entire range of phenotypic space. An alternative, more reasonable possibility is that the traits are not neutral, but rather are subject to some degree of purifying selection, which probably manifests itself as stabilizing selection, either real or apparent (Kondrashov and Turelli 1992). If so, the observed positive relationship between VG and VM is predicted at MSB. In an infinite population at MSB, 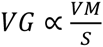, where *S* is the mean strength of selection against a new mutation; the proportionality becomes equality if the average selective effects are assumed to be uniform (Bulmer 1989; Barton 1990). If the average selective effects are not uniform (and surely they are not), 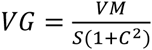 where *C* is the coefficient of variation of mutational effects on fitness (Charlesworth 2015), but unless *C* is highly variable among traits, VM/VG provides a reasonable approximation of the relative strength of selection. Thus, if genetic variation is maintained by MSB, unless the average strength of selection is very different between traits and/or the CVs of the mutational effects are very different, we expect a positive correlation between VG and VM. For example, if VM varies by two orders of magnitude, as it does here, selection would have to vary by nearly that much in order to remove the relationship between VG and VM.

The very strong relationship between VG and VM implies that, perhaps with a couple of exceptions, selection must be remarkably uniform across this set of traits. Why might that be? One possibility is that the traits themselves are all highly correlated, in which case direct selection on one trait might lead to sufficient indirect selection on the other traits to produce the pattern. There are too few degrees of freedom to calculate the full set of quadratic selection gradients for these traits (Lande and Arnold 1983), but a previous analysis of a subset of six of these traits (Traits 1, 2, 3, 15, 17, and 19) revealed that stabilizing selection on Embryo size (Trait 15) of strength V_s_ = VM/(VG)^2^ is sufficient to explain the observed standing genetic covariance matrix **G** for those traits, with no need to invoke selection on the other traits (Farhadifar *et al*. 2015). In that study, Embryo size was chosen *a priori* as the likely target of stabilizing selection, for three reasons. First, because from direct measurement of fecundity, we observed that embryo size showed the largest association with fecundity. Second, studies with many organisms have demonstrated that embryo size and size at birth are subject to stabilizing selection. And third, because it showed the largest deviation from the neutral expectation. The results of this study reinforce the previous finding: *t_P_* of Embryo size is about half that of the next smallest *t_P_* of the other 18 traits.

To some extent, the argument of the preceding paragraph begs the question, because all traits must be correlated with something, and one can never be certain one has accounted for all the relevant variables. Given the strong positive correlation between VM and VG for this particular set of traits, we can ask: where do other traits fall out in VM-VG space? Might it be that *any* arbitrary trait falls out more or less on the same line, and if so, why?

VG and VM have been previously quantified for four other traits in *C. elegans:* Lifetime reproduction weighted by survival measured under the MA conditions (W20), Lifetime reproduction measured in a high-throughput “worm-sorter” assay *(W_SORT_)*, median lifespan when exposed to the pathogenic bacteria *pseudomonas aeruginosa (LT50pa)* and body volume at maturity *(Size)* (Figure 3; Supplementary Table S5; Etienne *et al*. (2015)). Of the four traits, VG for *W_SORT_* and *LT50pa* were measured on nearly the same set of wild isolates as those reported in this study, so the values of VG and *t_P_* are directly comparable with those reported here. Persistence time for *W_SORT_* (166 generations) is almost identical to that of Embryo size (163 generations), and *t_P_* of *LT50pa* (335 generations) is on the low end of the spindle trait values. Persistence times for *W20* and *Size* are substantially smaller, but VG for those traits was measured on a smaller subset of wild isolates, some of which may be effectively the same clonal isolate. Unfortunately, only 11 isolates are common to the two datasets; for those 11 isolates VG and *t_P_* for *W_SORT_, W20*, and *Size* are both more similar to each other and closer to the common line, but the confidence limits are so large as to make the interpretation tenuous if not meaningless.

It is certainly within the realm of possibility that *t_P_* for more or less any trait measured in this set of wild isolates falls within the relatively narrow range observed here. We can think of at least two possible reasons why that might be. First, since *C. elegans* apparently experienced at least one hard, global, more or less genome-wide selective sweep within the recent past (~600-1250 generations; Andersen *et al*. (2012)), selection at linked loci must necessarily have been very inefficient immediately following the sweep, in which case the standing genetic variation may mostly represent a few hundred generations of input of effectively neutral mutations. The average persistence time of ~500 generations is consistent with that scenario. However, the two traits most clearly under selection on *a priori* grounds - Embryo size and lifetime reproduction - fall farthest below the line, which suggests, unsurprisingly, that some mutations are sufficiently deleterious as to have been effectively purged by selection.

A second possibility is that the predominantly self-fertilizing life history of *C. elegans*, combined with relatively restricted recombination within gene-rich regions of the genome (Rockman and Kruglyak 2009) means that most traits experience approximately the same overall level of background selection, although again, certain traits clearly experience atypically strong (or weak) selection.

The mean-standardized VM varies over nearly two orders of magnitude (~80X) for the traits reported in this study. Two classes of (non-exclusive) explanations for variation in VM among traits have been put forth. First, for whatever reason, the different traits may be differently robust to the effects of mutation, or in other words, for a given number of mutations that affect a trait, the average effect on the phenotype differs among traits. Stearns and Kawecki (1994) proposed that VM provides a measure of the robustness of a trait to the perturbing effects of new mutations, such that 1/VM is a meaningful estimate of mutational robustness. Alternatively, the traits may present different mutational targets, i.e. different numbers of genetic loci may potentially affect the trait. Presumably, fewer loci affect (say) the expression level of Gene X than affect fitness. However, Houle (1998) pointed out that VM cannot provide an unambiguous measure of mutational robustness because of the confounding influence of target size. Moreover, there is some conceptual ambiguity between target size and effect size, because it is logically coherent to say that all traits have the same target - the genome - but the number of loci whose effect is zero differs between traits.

For these traits, the average effect (|*α*|) clearly differs in the expected way, i.e., in 3/3 cases |*α*| is greater in the high VM trait than in the low VM trait. If the mutational target also differs in the expected way, the best-fit *UQ_z_* for high-VM traits should be greater than that for low-VM traits, and that is not true in any of the cases we examined.

Houle’s critique of Stearns’ and Kawecki’s Genetic Robustness argument has been as influential as it is logically convincing: VM is no longer considered a reliable measure of mutational robustness (e.g., Gibson and Wagner 2000). We find ourselves in the unexpected position of suggesting that VM may deserve a second look as a potentially meaningful measure of mutational robustness.

### Conclusions and Future Directions

1. For this set of 19 functionally related traits, the mutational process is very repeatable. There are several other organisms for which there are extant MA lines from multiple starting genotypes (e.g., *Chlamydomonas reinhardtii*, (Ness *et al*. 2015); *Caenorhabditis briggsae* and *Oscheius myriophila* (Baer *et al*. 2005), *Arabidopsis thaliana* (C. Fenster, personal communication); *Daphnia pulex* (S. Schaack, personal communication) and probably others. Similar studies including suites of different types of traits will help establish the boundaries of repeatability and idiosyncrasy in the mutational process.

2. For these traits in this species, mean-standardized VM almost perfectly predicts VG. This result has been previously documented, in *D. melanogaster* (Houle 1998) and to a lesser extent in *D. pulex* (reanalysis of data in Lynch *et al*. 1998, above). It is the predicted result if genetic variation is predominantly due to mutation-selection balance or the interplay between mutation and drift. It is not predicted if balancing selection of primary importance in the maintenance of genetic variation.

*C. elegans* is, *prima facie*, an unlikely target for balancing selection because of the strong evidence for a recent episode of global strong directional selection (Andersen *et al*. 2012). However, recent evidence suggests that balancing selection may have maintained variation in numerous regions throughout the *C. elegans* genome (Thompson *et al*. 2015). Looking farther afield, it has been very convincingly argued that VM is not a sufficient predictor of VG for life-history traits in Drosophila, and that there must be a significant contribution to VG from balancing selection (Charlesworth 2015). If so, the effect of balancing selection would effectively be to move the line of relationship between VG and VM (depicted in Figure 3) upwards, i.e. to increase the intercept. If balancing selection contributes more to VG for life history traits than for other classes of traits, it implies that, all else equal, the slope of the relationship between VG and VM will be steeper than the line of neutrality, with persistence times of high-VM life history traits falling closer to the line of neutrality. All else is not equal, however; persistence times for life history traits in *D. melanogaster* and other taxa are, on average, less than half those for morphological or other traits (Houle *et al*. 1996; Houle 1998; Lynch *et al*. 1999).

3. Whether differences in VM between traits are better explained by differences in target size or mutational robustness (effect size) cannot be conclusively determined from these data, although to the extent that our simple models reflect reality, it appears that differences in VM are better explained by differences in robustness. To more satisfactorily address this important issue, it will be useful to compare sets of traits that differ substantially in both VM and in mutational heritability. Because of the strong positive relationship between VM and VE, this may be easier said than done in most cases, although the outliers may be particularly informative.

## Acknowledgments

We thank Asher Cutter, Dee Denver, Marie-Anne Félix, Karin Kiontke, Ralf Sommer, and the Caenorhabditis Genetics Center (CGC) for providing wild isolates. The CGC is funded by the NIH Office of Research Infrastructure Programs (P40 OD010440). Support was provided by Human Frontier Science Program grant RGP 0034/2010 to T. Müller-Reichert, M. Delattre, and DJN, NIH award R01GM072639 to CFB and D. R. Denver, and NIH award R01GM107227 to CFB, ECA and JMP.

---

**Supplementary Figure S1.**
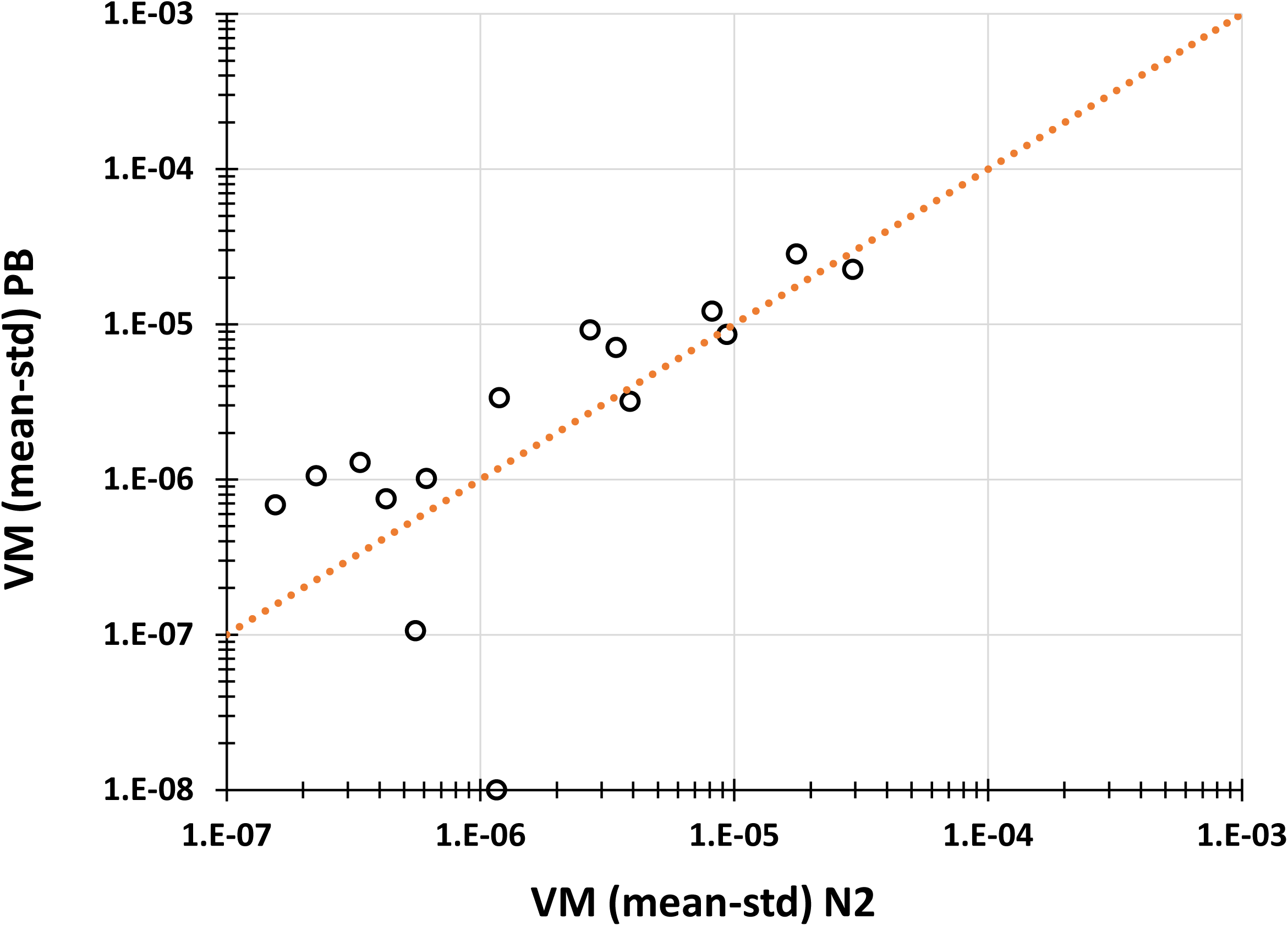
Mean-standardized VM (PB306) plotted against mean-standardized VM (N2). The dashed line represents the line of equality.

**Supplementary Figure S2.**
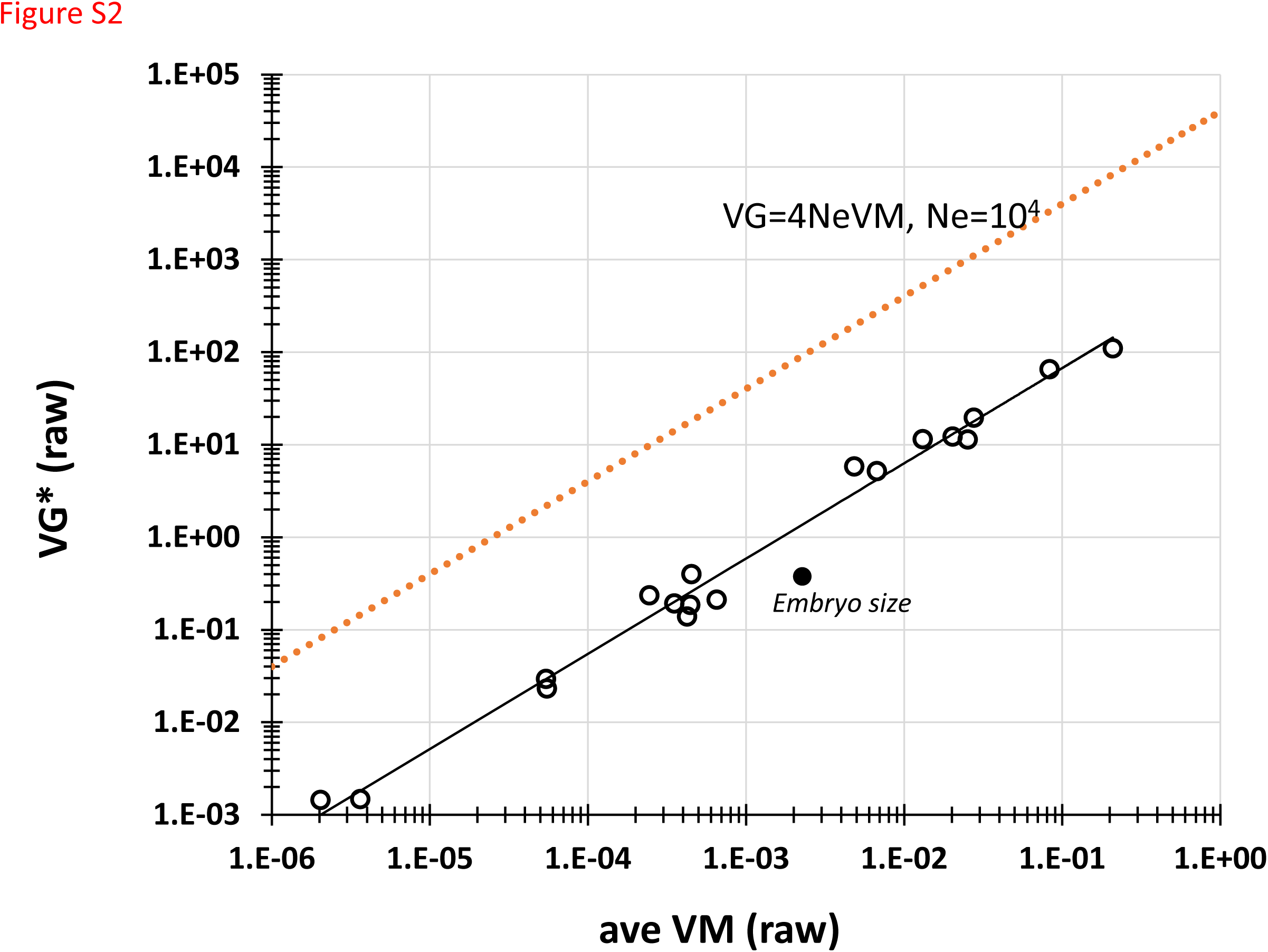
Raw VG^*^ plotted against raw VM. The solid black line shows the best-fit of the spindle trait data. The orange dashed line shows 4*N_e_*VM for *N_e_* = 10^4^.

**Table S2.A.**
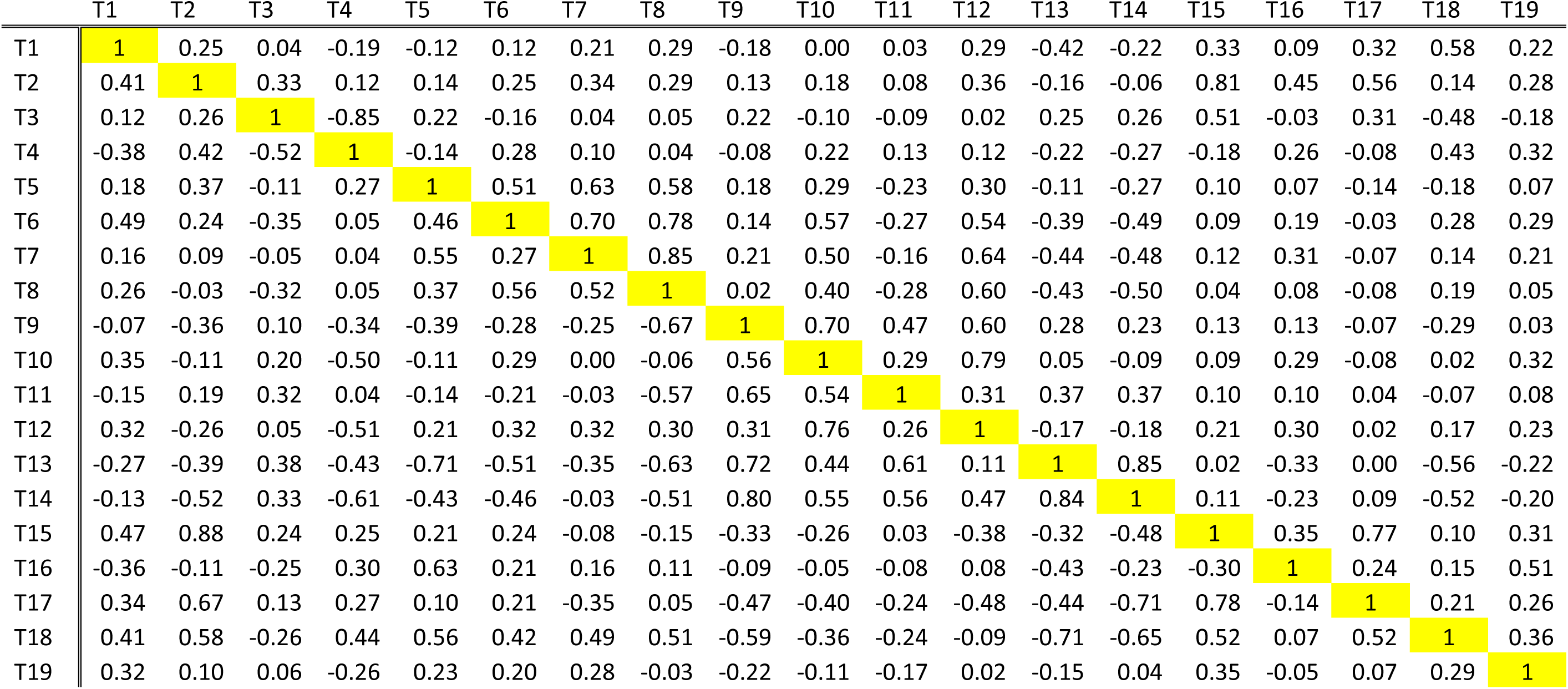
Pairwise correlations of N2 line means. MA lines above the diagonal, G0 pseudolines below the diagonal.

|  | T1 | T2 | T3 | T4 | T5 | T6 | T7 | T8 | T9 | T10 | T11 | T12 | T13 | T14 | T15 | T16 | T17 | T18 | T19 |
| --- | --- | --- | --- | --- | --- | --- | --- | --- | --- | --- | --- | --- | --- | --- | --- | --- | --- | --- | --- |
| T1 | 1 | 0.25 | 0.04 | -0.19 | -0.12 | 0.12 | 0.21 | 0.29 | -0.18 | 0.00 | 0.03 | 0.29 | -0.42 | -0.22 | 0.33 | 0.09 | 0.32 | 0.58 | 0.22 |
| T2 | 0.41 | 1 | 0.33 | 0.12 | 0.14 | 0.25 | 0.34 | 0.29 | 0.13 | 0.18 | 0.08 | 0.36 | -0.16 | -0.06 | 0.81 | 0.45 | 0.56 | 0.14 | 0.28 |
| T3 | 0.12 | 0.26 | 1 | -0.85 | 0.22 | -0.16 | 0.04 | 0.05 | 0.22 | -0.10 | -0.09 | 0.02 | 0.25 | 0.26 | 0.51 | -0.03 | 0.31 | -0.48 | -0.18 |
| T4 | -0.38 | 0.42 | -0.52 | 1 | -0.14 | 0.28 | 0.10 | 0.04 | -0.08 | 0.22 | 0.13 | 0.12 | -0.22 | -0.27 | -0.18 | 0.26 | -0.08 | 0.43 | 0.32 |
| T5 | 0.18 | 0.37 | -0.11 | 0.27 | 1 | 0.51 | 0.63 | 0.58 | 0.18 | 0.29 | -0.23 | 0.30 | -0.11 | -0.27 | 0.10 | 0.07 | -0.14 | -0.18 | 0.07 |
| T6 | 0.49 | 0.24 | -0.35 | 0.05 | 0.46 | 1 | 0.70 | 0.78 | 0.14 | 0.57 | -0.27 | 0.54 | -0.39 | -0.49 | 0.09 | 0.19 | -0.03 | 0.28 | 0.29 |
| T7 | 0.16 | 0.09 | -0.05 | 0.04 | 0.55 | 0.27 | 1 | 0.85 | 0.21 | 0.50 | -0.16 | 0.64 | -0.44 | -0.48 | 0.12 | 0.31 | -0.07 | 0.14 | 0.21 |
| T8 | 0.26 | -0.03 | -0.32 | 0.05 | 0.37 | 0.56 | 0.52 | 1 | 0.02 | 0.40 | -0.28 | 0.60 | -0.43 | -0.50 | 0.04 | 0.08 | -0.08 | 0.19 | 0.05 |
| T9 | -0.07 | -0.36 | 0.10 | -0.34 | -0.39 | -0.28 | -0.25 | -0.67 | 1 | 0.70 | 0.47 | 0.60 | 0.28 | 0.23 | 0.13 | 0.13 | -0.07 | -0.29 | 0.03 |
| T10 | 0.35 | -0.11 | 0.20 | -0.50 | -0.11 | 0.29 | 0.00 | -0.06 | 0.56 | 1 | 0.29 | 0.79 | 0.05 | -0.09 | 0.09 | 0.29 | -0.08 | 0.02 | 0.32 |
| T11 | -0.15 | 0.19 | 0.32 | 0.04 | -0.14 | -0.21 | -0.03 | -0.57 | 0.65 | 0.54 | 1 | 0.31 | 0.37 | 0.37 | 0.10 | 0.10 | 0.04 | -0.07 | 0.08 |
| T12 | 0.32 | -0.26 | 0.05 | -0.51 | 0.21 | 0.32 | 0.32 | 0.30 | 0.31 | 0.76 | 0.26 | 1 | -0.17 | -0.18 | 0.21 | 0.30 | 0.02 | 0.17 | 0.23 |
| T13 | -0.27 | -0.39 | 0.38 | -0.43 | -0.71 | -0.51 | -0.35 | -0.63 | 0.72 | 0.44 | 0.61 | 0.11 | 1 | 0.85 | 0.02 | -0.33 | 0.00 | -0.56 | -0.22 |
| T14 | -0.13 | -0.52 | 0.33 | -0.61 | -0.43 | -0.46 | -0.03 | -0.51 | 0.80 | 0.55 | 0.56 | 0.47 | 0.84 | 1 | 0.11 | -0.23 | 0.09 | -0.52 | -0.20 |
| T15 | 0.47 | 0.88 | 0.24 | 0.25 | 0.21 | 0.24 | -0.08 | -0.15 | -0.33 | -0.26 | 0.03 | -0.38 | -0.32 | -0.48 | 1 | 0.35 | 0.77 | 0.10 | 0.31 |
| T16 | -0.36 | -0.11 | -0.25 | 0.30 | 0.63 | 0.21 | 0.16 | 0.11 | -0.09 | -0.05 | -0.08 | 0.08 | -0.43 | -0.23 | -0.30 | 1 | 0.24 | 0.15 | 0.51 |
| T17 | 0.34 | 0.67 | 0.13 | 0.27 | 0.10 | 0.21 | -0.35 | 0.05 | -0.47 | -0.40 | -0.24 | -0.48 | -0.44 | -0.71 | 0.78 | -0.14 | 1 | 0.21 | 0.26 |
| T18 | 0.41 | 0.58 | -0.26 | 0.44 | 0.56 | 0.42 | 0.49 | 0.51 | -0.59 | -0.36 | -0.24 | -0.09 | -0.71 | -0.65 | 0.52 | 0.07 | 0.52 | 1 | 0.36 |
| T19 | 0.32 | 0.10 | 0.06 | -0.26 | 0.23 | 0.20 | 0.28 | -0.03 | -0.22 | -0.11 | -0.17 | 0.02 | -0.15 | 0.04 | 0.35 | -0.05 | 0.07 | 0.29 | 1 |

**Table S2.B.** Pairwise correlations of PB306 line means. MA lines above the diagonal, G0 pseudolines below the diagonal.

|  | T1 | T2 | T3 | T4 | T5 | T6 | T7 | T8 | T9 | T10 | T11 | T12 | T13 | T14 | T15 | T16 | T17 | T18 | T19 |
| --- | --- | --- | --- | --- | --- | --- | --- | --- | --- | --- | --- | --- | --- | --- | --- | --- | --- | --- | --- |
| T1 | 1 | 0.24 | -0.01 | -0.21 | -0.10 | -0.08 | 0.05 | 0.01 | 0.13 | 0.23 | 0.33 | 0.26 | 0.01 | -0.16 | 0.19 | -0.06 | 0.27 | 0.52 | 0.15 |
| T2 | 0.41 | 1 | 0.42 | 0.19 | 0.06 | 0.41 | 0.01 | 0.35 | 0.00 | 0.31 | -0.29 | 0.20 | 0.00 | -0.07 | 0.81 | -0.06 | 0.77 | 0.36 | 0.70 |
| T3 | 0.12 | 0.26 | 1 | -0.72 | 0.09 | -0.09 | -0.14 | -0.14 | 0.35 | 0.04 | 0.07 | 0.05 | -0.06 | 0.22 | 0.67 | 0.05 | 0.64 | -0.35 | 0.09 |
| T4 | -0.38 | 0.42 | -0.52 | 1 | -0.04 | 0.48 | 0.14 | 0.42 | -0.48 | 0.14 | -0.46 | 0.00 | 0.09 | -0.24 | -0.22 | -0.07 | -0.22 | 0.41 | 0.42 |
| T5 | 0.18 | 0.37 | -0.11 | 0.27 | 1 | 0.35 | 0.38 | 0.44 | 0.07 | 0.16 | -0.22 | 0.04 | -0.45 | -0.29 | 0.00 | 0.07 | -0.04 | -0.13 | 0.19 |
| T6 | 0.49 | 0.24 | -0.35 | 0.05 | 0.46 | 1 | 0.48 | 0.81 | -0.46 | 0.54 | -0.70 | 0.23 | -0.05 | -0.41 | 0.10 | -0.22 | 0.11 | 0.26 | 0.54 |
| T7 | 0.16 | 0.09 | -0.05 | 0.04 | 0.55 | 0.27 | 1 | 0.79 | 0.07 | 0.52 | -0.16 | 0.59 | 0.00 | -0.22 | -0.27 | -0.13 | -0.25 | 0.09 | 0.04 |
| T8 | 0.26 | -0.03 | -0.32 | 0.05 | 0.37 | 0.56 | 0.52 | 1 | -0.14 | 0.58 | -0.49 | 0.55 | 0.01 | -0.27 | -0.04 | -0.23 | -0.01 | 0.27 | 0.36 |
| T9 | -0.07 | -0.36 | 0.10 | -0.34 | -0.39 | -0.28 | -0.25 | -0.67 | 1 | 0.27 | 0.68 | 0.46 | 0.25 | 0.45 | 0.15 | 0.03 | 0.06 | -0.24 | -0.36 |
| T10 | 0.35 | -0.11 | 0.20 | -0.50 | -0.11 | 0.29 | 0.00 | -0.06 | 0.56 | 1 | -0.01 | 0.70 | 0.29 | -0.15 | 0.10 | -0.19 | 0.07 | 0.18 | 0.23 |
| T11 | -0.15 | 0.19 | 0.32 | 0.04 | -0.14 | -0.21 | -0.03 | -0.57 | 0.65 | 0.54 | 1 | 0.27 | 0.13 | 0.28 | -0.02 | 0.20 | -0.06 | -0.17 | -0.52 |
| T12 | 0.32 | -0.26 | 0.05 | -0.51 | 0.21 | 0.32 | 0.32 | 0.30 | 0.31 | 0.76 | 0.26 | 1 | 0.37 | 0.08 | 0.03 | -0.12 | -0.01 | 0.17 | 0.00 |
| T13 | -0.27 | -0.39 | 0.38 | -0.43 | -0.71 | -0.51 | -0.35 | -0.63 | 0.72 | 0.44 | 0.61 | 0.11 | 1 | 0.70 | -0.11 | -0.46 | -0.15 | -0.12 | -0.13 |
| T14 | -0.13 | -0.52 | 0.33 | -0.61 | -0.43 | -0.46 | -0.03 | -0.51 | 0.80 | 0.55 | 0.56 | 0.47 | 0.84 | 1 | -0.01 | -0.35 | -0.05 | -0.50 | -0.28 |
| T15 | 0.47 | 0.88 | 0.24 | 0.25 | 0.21 | 0.24 | -0.08 | -0.15 | -0.33 | -0.26 | 0.03 | -0.38 | -0.32 | -0.48 | 1 | 0.06 | 0.88 | 0.11 | 0.42 |
| T16 | -0.36 | -0.11 | -0.25 | 0.30 | 0.63 | 0.21 | 0.16 | 0.11 | -0.09 | -0.05 | -0.08 | 0.08 | -0.43 | -0.23 | -0.30 | 1 | 0.05 | 0.06 | 0.01 |
| T17 | 0.34 | 0.67 | 0.13 | 0.27 | 0.10 | 0.21 | -0.35 | 0.05 | -0.47 | -0.40 | -0.24 | -0.48 | -0.44 | -0.71 | 0.78 | -0.14 | 1 | 0.22 | 0.49 |
| T18 | 0.41 | 0.58 | -0.26 | 0.44 | 0.56 | 0.42 | 0.49 | 0.51 | -0.59 | -0.36 | -0.24 | -0.09 | -0.71 | -0.65 | 0.52 | 0.07 | 0.52 | 1 | 0.44 |
| T19 | 0.32 | 0.10 | 0.06 | -0.26 | 0.23 | 0.20 | 0.28 | -0.03 | -0.22 | -0.11 | -0.17 | 0.02 | -0.15 | 0.04 | 0.35 | -0.05 | 0.07 | 0.29 | 1 |

**Table S2.C.** Correlations of wild isolate line means.

|  | T1 | T2 | T3 | T4 | T5 | T6 | T7 | T8 | T9 | T10 | T11 | T12 | T13 | T14 | T15 | T16 | T17 | T18 | T19 |
| --- | --- | --- | --- | --- | --- | --- | --- | --- | --- | --- | --- | --- | --- | --- | --- | --- | --- | --- | --- |
| T1 | 0 | 0.13 | -0.09 | 0.20 | -0.36 | -0.08 | 0.02 | 0.08 | 0.18 | 0.07 | 0.48 | 0.24 | 0.03 | -0.10 | 0.05 | 0.25 | 0.02 | 0.33 | -0.11 |
| T2 |  | 0 | 0.38 | -0.41 | -0.23 | -0.18 | -0.12 | 0.03 | 0.07 | -0.02 | -0.21 | 0.16 | 0.24 | 0.36 | 0.00 | 0.27 | 0.32 | -0.40 | 0.18 |
| T3 |  |  | 0 | -0.12 | 0.21 | 0.22 | 0.05 | 0.23 | 0.13 | -0.14 | -0.23 | -0.09 | -0.32 | -0.15 | 0.47 | 0.25 | 0.53 | -0.12 | 0.12 |
| T4 |  |  |  | 0 | -0.30 | 0.02 | -0.11 | -0.05 | -0.20 | 0.22 | -0.21 | 0.19 | 0.31 | 0.26 | -0.51 | -0.13 | -0.32 | -0.08 | 0.33 |
| T5 |  |  |  |  | 0 | 0.03 | -0.15 | 0.10 | 0.09 | 0.02 | -0.19 | -0.22 | 0.19 | 0.04 | -0.12 | -0.18 | -0.08 | -0.56 | -0.01 |
| T6 |  |  |  |  |  | 0 | 0.26 | 0.16 | -0.14 | 0.12 | -0.34 | 0.04 | 0.24 | 0.02 | -0.27 | -0.16 | -0.08 | -0.25 | 0.09 |
| T7 |  |  |  |  |  |  | 0 | 0.19 | 0.05 | 0.30 | -0.15 | 0.11 | 0.10 | -0.15 | -0.13 | 0.04 | 0.20 | -0.31 | -0.15 |
| T8 |  |  |  |  |  |  |  | 0 | 0.10 | 0.32 | -0.04 | 0.13 | 0.24 | 0.07 | -0.07 | -0.14 | 0.05 | -0.29 | -0.04 |
| T9 |  |  |  |  |  |  |  |  | 0 | -0.21 | -0.11 | -0.05 | -0.32 | -0.27 | 0.11 | 0.08 | 0.28 | 0.08 | 0.05 |
| T10 |  |  |  |  |  |  |  |  |  | 0 | -0.38 | -0.01 | -0.19 | -0.49 | -0.11 | 0.07 | 0.11 | 0.18 | 0.36 |
| T11 |  |  |  |  |  |  |  |  |  |  | 0 | -0.05 | -0.31 | -0.25 | 0.01 | 0.20 | 0.18 | 0.11 | 0.09 |
| T12 |  |  |  |  |  |  |  |  |  |  |  | 0 | -0.09 | -0.34 | 0.09 | 0.09 | 0.39 | 0.15 | 0.16 |
| T13 |  |  |  |  |  |  |  |  |  |  |  |  | 0 | -0.08 | 0.07 | -0.12 | 0.07 | 0.27 | 0.20 |
| T14 |  |  |  |  |  |  |  |  |  |  |  |  |  | 0 | 0.30 | -0.13 | 0.32 | 0.04 | 0.04 |
| T15 |  |  |  |  |  |  |  |  |  |  |  |  |  |  | 0 | 0.23 | 0.20 | -0.30 | 0.04 |
| T16 |  |  |  |  |  |  |  |  |  |  |  |  |  |  |  | 0 | 0.11 | 0.10 | 0.15 |
| T17 |  |  |  |  |  |  |  |  |  |  |  |  |  |  |  |  | 0 | 0.07 | 0.29 |
| T18 |  |  |  |  |  |  |  |  |  |  |  |  |  |  |  |  |  | 0 | 0.00 |
| T19 |  |  |  |  |  |  |  |  |  |  |  |  |  |  |  |  |  |  | 0 |

**Table S3.** Simulation parameters. Column headings: *Trait/Strain*, Trait number/strain; *DME*, distribution of effects model (Exponential or Normal); *UQ_z_*, number of mutations per-generation affecting the trait; *α^2^*, average squared mutational effect, scaled as a fraction of the G0 mean; |*α*|,absolute value of the mutational effect; *VM (=Uα^2^)*, mean-standardized mutational variance; *VE^*^*, environmental variance (see text for details); *E(#muts/line)*, expected number of mutations per MA line; *nbest/1000;* proportion of simulations in which the model was best; *E[distance];* average KL distance of the simulated data from observed.

| *Trait/Strain* | *DME* | *UQ<sub>z</sub>* | $\alpha^2$ | $\vert \alpha\vert $ | *VM (=UQ<sub>z</sub>$\alpha^2$)* | *VE\** | *nbest/1000* | *E[Distance]* |
| --- | --- | --- | --- | --- | --- | --- | --- | --- |
| *8 / pool* | Exp | $5 \times 10^{-1}$ | $4.6 \times 10^{-5}$ | 0.0068 | $2.30 \times 10^{-5}$ | 0.107 | 43 | 25.01 |
| *8 / pool* | Exp | $1 \times 10^{-1}$ | $2.3 \times 10^{-4}$ | 0.015 | $2.30 \times 10^{-5}$ | 0.107 | 57 | 25.01 |
| *8 / pool* | Exp | $2 \times 10^{-2}$ | $1.2 \times 10^{-3}$ | 0.034 | $2.30 \times 10^{-5}$ | 0.107 | 68 | 24.80 |
| *8 / pool* | Exp | $4 \times 10^{-3}$ | $5.8 \times 10^{-3}$ | 0.076 | $2.30 \times 10^{-5}$ | 0.107 | 142 | 24.32 |
| *8 / pool* | Exp | $8 \times 10^{-4}$ | $2.9 \times 10^{-2}$ | 0.17 | $2.30 \times 10^{-5}$ | 0.107 | 141 | 24.60 |
| *8 / pool* | Exp | $1.6 \times 10^{-4}$ | $1.4 \times 10^{-1}$ | 0.38 | $2.30 \times 10^{-5}$ | 0.107 | 16 | 27.20 |
| *8 / pool* | Exp | $3.2 \times 10^{-5}$ | $7.2 \times 10^{-1}$ | 0.85 | $2.30 \times 10^{-5}$ | 0.107 | 0 | 29.31 |
| *8 / pool* | Exp | $6.4 \times 10^{-6}$ | 3.6 | 1.90 | $2.30 \times 10^{-5}$ | 0.107 | 0 | 29.92 |
| *8 / pool* | Norm | $5 \times 10^{-1}$ | $4.6 \times 10^{-5}$ | 0.0068 | $2.30 \times 10^{-5}$ | 0.107 | 45 | 25.01 |
| *8 / pool* | Norm | $1 \times 10^{-1}$ | $2.3 \times 10^{-4}$ | 0.015 | $2.30 \times 10^{-5}$ | 0.107 | 40 | 25.02 |
| *8 / pool* | Norm | $2 \times 10^{-2}$ | $1.2 \times 10^{-3}$ | 0.034 | $2.30 \times 10^{-5}$ | 0.107 | 71 | 24.86 |
| *8 / pool* | Norm | $4 \times 10^{-3}$ | $5.8 \times 10^{-3}$ | 0.076 | $2.30 \times 10^{-5}$ | 0.107 | 104 | 24.46 |
| *8 / pool* | Norm | $8 \times 10^{-4}$ | $2.9 \times 10^{-2}$ | 0.17 | $2.30 \times 10^{-5}$ | 0.107 | 240 | 23.37 |
| *8 / pool* | Norm | $1.6 \times 10^{-4}$ | $1.4 \times 10^{-1}$ | 0.38 | $2.30 \times 10^{-5}$ | 0.107 | 33 | 26.73 |
| *8 / pool* | Norm | $3.2 \times 10^{-5}$ | $7.2 \times 10^{-1}$ | 0.85 | $2.30 \times 10^{-5}$ | 0.107 | 0 | 29.13 |
| *8 / pool* | Norm | $6.4 \times 10^{-6}$ | 3.6 | 1.90 | $2.30 \times 10^{-5}$ | 0.107 | 0 | 29.99 |

| *Trait/Strain* | *DME* | *UQ<sub>z</sub>* | *α<sup>2</sup>* | *\|α\|* | *VM (=UQ<sub>z</sub>α<sup>2</sup>)* | *VE\** | *nbest/1000* | *E[Distance]* |
| --- | --- | --- | --- | --- | --- | --- | --- | --- |
| 17 / pool | Exp | $5 \times 10^{-1}$ | $1.6 \times 10^{-6}$ | 0.0013 | $8.14 \times 10^{-7}$ | 0.0046 | 55 | 24.98 |
| 17 / pool | Exp | $1 \times 10^{-1}$ | $8.1 \times 10^{-6}$ | 0.0029 | $8.14 \times 10^{-7}$ | 0.0046 | 62 | 24.96 |
| 17 / pool | Exp | $2 \times 10^{-2}$ | $4.1 \times 10^{-5}$ | 0.0064 | $8.14 \times 10^{-7}$ | 0.0046 | 68 | 24.79 |
| 17 / pool | Exp | $4 \times 10^{-3}$ | $2.0 \times 10^{-4}$ | 0.14 | $8.14 \times 10^{-7}$ | 0.0046 | 171 | 24.16 |
| 17 / pool | Exp | $8 \times 10^{-4}$ | $1.0 \times 10^{-3}$ | 0.32 | $8.14 \times 10^{-7}$ | 0.0046 | 158 | 24.56 |
| 17 / pool | Exp | $1.6 \times 10^{-4}$ | $5.1 \times 10^{-3}$ | 0.071 | $8.14 \times 10^{-7}$ | 0.0046 | 13 | 27.33 |
| 17 / pool | Exp | $3.2 \times 10^{-5}$ | $2.5 \times 10^{-2}$ | 0.16 | $8.14 \times 10^{-7}$ | 0.0046 | 0 | 29.25 |
| 17 / pool | Exp | $6.4 \times 10^{-6}$ | $1.3 \times 10^{-1}$ | 0.36 | $8.14 \times 10^{-7}$ | 0.0046 | 0 | 29.94 |
| 17 / pool | Norm | $5 \times 10^{-1}$ | $1.6 \times 10^{-6}$ | 0.0013 | $8.14 \times 10^{-7}$ | 0.0046 | 9 | 25.05 |
| 17 / pool | Norm | $1 \times 10^{-1}$ | $8.1 \times 10^{-6}$ | 0.0029 | $8.14 \times 10^{-7}$ | 0.0046 | 54 | 25.03 |
| 17 / pool | Norm | $2 \times 10^{-2}$ | $4.1 \times 10^{-5}$ | 0.0064 | $8.14 \times 10^{-7}$ | 0.0046 | 52 | 24.93 |
| 17 / pool | Norm | $4 \times 10^{-3}$ | $2.0 \times 10^{-4}$ | 0.14 | $8.14 \times 10^{-7}$ | 0.0046 | 101 | 24.42 |
| 17 / pool | Norm | $8 \times 10^{-4}$ | $1.0 \times 10^{-3}$ | 0.32 | $8.14 \times 10^{-7}$ | 0.0046 | 173 | 24.10 |
| 17 / pool | Norm | $1.6 \times 10^{-4}$ | $5.1 \times 10^{-3}$ | 0.071 | $8.14 \times 10^{-7}$ | 0.0046 | 36 | 26.68 |
| 17 / pool | Norm | $3.2 \times 10^{-5}$ | $2.5 \times 10^{-2}$ | 0.16 | $8.14 \times 10^{-7}$ | 0.0046 | 1 | 29.22 |
| 17 / pool | Norm | $6.4 \times 10^{-6}$ | $1.3 \times 10^{-1}$ | 0.36 | $8.14 \times 10^{-7}$ | 0.0046 | 0 | 29.94 |

| *Trait/Strain* | *DME* | *UQ<sub>z</sub>* | $\alpha^2$ | $\vert \alpha\vert $ | *VM (=UQ<sub>z</sub>$\alpha^2$)* | *VE\** | *nbest/1000* | *E(distance)* |
| --- | --- | --- | --- | --- | --- | --- | --- | --- |
| 7/N2 | Exp | $5 \times 10^{-1}$ | $5.9 \times 10^{-5}$ | 0.0077 | $2.94 \times 10^{-5}$ | 0.088 | 123 | 23.21 |
| 7/N2 | Exp | $1 \times 10^{-1}$ | $2.9 \times 10^{-4}$ | 0.017 | $2.94 \times 10^{-5}$ | 0.088 | 144 | 23.32 |
| 7/N2 | Exp | $2 \times 10^{-2}$ | $1.5 \times 10^{-3}$ | 0.038 | $2.94 \times 10^{-5}$ | 0.088 | 113 | 23.38 |
| 7/N2 | Exp | $4 \times 10^{-3}$ | $7.3 \times 10^{-3}$ | 0.086 | $2.94 \times 10^{-5}$ | 0.088 | 93 | 23.99 |
| 7/N2 | Exp | $8 \times 10^{-4}$ | $3.7 \times 10^{-2}$ | 0.19 | $2.94 \times 10^{-5}$ | 0.088 | 15 | 26.36 |
| 7/N2 | Exp | $1.6 \times 10^{-4}$ | $1.8 \times 10^{-1}$ | 0.43 | $2.94 \times 10^{-5}$ | 0.088 | 0 | 29.04 |
| 7/N2 | Exp | $3.2 \times 10^{-5}$ | $9.2 \times 10^{-1}$ | 0.96 | $2.94 \times 10^{-5}$ | 0.088 | 0 | 30.35 |
| 7/N2 | Exp | $6.4 \times 10^{-6}$ | 4.6 | 2.14 | $2.94 \times 10^{-5}$ | 0.088 | 0 | 30.80 |
| 7/N2 | Norm | $5 \times 10^{-1}$ | $5.9 \times 10^{-5}$ | 0.0077 | $2.94 \times 10^{-5}$ | 0.088 | 147 | 23.25 |
| 7/N2 | Norm | $1 \times 10^{-1}$ | $2.9 \times 10^{-4}$ | 0.017 | $2.94 \times 10^{-5}$ | 0.088 | 113 | 23.25 |
| 7/N2 | Norm | $2 \times 10^{-2}$ | $1.5 \times 10^{-3}$ | 0.038 | $2.94 \times 10^{-5}$ | 0.088 | 110 | 23.34 |
| 7/N2 | Norm | $4 \times 10^{-3}$ | $7.3 \times 10^{-3}$ | 0.086 | $2.94 \times 10^{-5}$ | 0.088 | 112 | 23.52 |
| 7/N2 | Norm | $8 \times 10^{-4}$ | $3.7 \times 10^{-2}$ | 0.19 | $2.94 \times 10^{-5}$ | 0.088 | 23 | 25.56 |
| 7/N2 | Norm | $1.6 \times 10^{-4}$ | $1.8 \times 10^{-1}$ | 0.43 | $2.94 \times 10^{-5}$ | 0.088 | 0 | 28.71 |
| 7/N2 | Norm | $3.2 \times 10^{-5}$ | $9.2 \times 10^{-1}$ | 0.96 | $2.94 \times 10^{-5}$ | 0.088 | 0 | 30.21 |
| 7/N2 | Norm | $6.4 \times 10^{-6}$ | 4.6 | 2.14 | $2.94 \times 10^{-5}$ | 0.088 | 0 | 30.60 |
| 7/PB | Exp | $5 \times 10^{-1}$ | $4.5 \times 10^{-5}$ | 0.0067 | $2.26 \times 10^{-5}$ | 0.098 | 10 | 16.95 |
| 7/PB | Exp | $1 \times 10^{-1}$ | $2.3 \times 10^{-4}$ | 0.015 | $2.26 \times 10^{-5}$ | 0.098 | 15 | 16.88 |
| 7/PB | Exp | $2 \times 10^{-2}$ | $1.1 \times 10^{-3}$ | 0.033 | $2.26 \times 10^{-5}$ | 0.098 | 13 | 16.79 |
| 7/PB | Exp | $4 \times 10^{-3}$ | $5.6 \times 10^{-3}$ | 0.075 | $2.26 \times 10^{-5}$ | 0.098 | 23 | 15.92 |
| 7/PB | Exp | $8 \times 10^{-4}$ | $2.8 \times 10^{-2}$ | 0.17 | $2.26 \times 10^{-5}$ | 0.098 | 73 | 14.82 |
| 7/PB | Exp | $1.6 \times 10^{-4}$ | $1.4 \times 10^{-2}$ | 0.38 | $2.26 \times 10^{-5}$ | 0.098 | 124 | 14.12 |
| 7/PB | Exp | $3.2 \times 10^{-5}$ | $0.70 \times 10^{-1}$ | 0.84 | $2.26 \times 10^{-5}$ | 0.098 | 124 | 14.10 |
| 7/PB | Exp | $6.4 \times 10^{-6}$ | 3.52 | 1.88 | $2.26 \times 10^{-5}$ | 0.098 | 149 | 14.02 |
| 7/PB | Norm | $5 \times 10^{-1}$ | $4.5 \times 10^{-5}$ | 0.0067 | $2.26 \times 10^{-5}$ | 0.098 | 11 | 16.87 |
| 7/PB | Norm | $1 \times 10^{-1}$ | $2.3 \times 10^{-4}$ | 0.015 | $2.26 \times 10^{-5}$ | 0.098 | 13 | 16.94 |
| 7/PB | Norm | $2 \times 10^{-2}$ | $1.1 \times 10^{-3}$ | 0.033 | $2.26 \times 10^{-5}$ | 0.098 | 12 | 16.91 |
| 7/PB | Norm | $4 \times 10^{-3}$ | $5.6 \times 10^{-3}$ | 0.075 | $2.26 \times 10^{-5}$ | 0.098 | 15 | 16.26 |
| 7/PB | Norm | $8 \times 10^{-4}$ | $2.8 \times 10^{-2}$ | 0.17 | $2.26 \times 10^{-5}$ | 0.098 | 65 | 15.06 |
| 7/PB | Norm | $1.6 \times 10^{-4}$ | $1.4 \times 10^{-2}$ | 0.38 | $2.26 \times 10^{-5}$ | 0.098 | 112 | 14.34 |
| 7/PB | Norm | $3.2 \times 10^{-5}$ | $0.70 \times 10^{-1}$ | 0.84 | $2.26 \times 10^{-5}$ | 0.098 | 128 | 14.10 |
| 7/PB | Norm | $6.4 \times 10^{-6}$ | 3.52 | 1.88 | $2.26 \times 10^{-5}$ | 0.098 | 113 | 13.97 |

| *Trait/Strain* | *DME* | *UQ<sub>z</sub>* | *α<sup>2</sup>* | *\|α\|* | *VM (=UQ<sub>z</sub>α<sup>2</sup>)* | *VE\** | *nbest/1000* | *E(distance)* |
| --- | --- | --- | --- | --- | --- | --- | --- | --- |
| 2/N2 | Exp | $5 \times 10^{-1}$ | $4.5 \times 10^{-7}$ | 0.00067 | $2.26 \times 10^{-7}$ | 0.0026 | 94 | 22.07 |
| 2/N2 | Exp | $1 \times 10^{-1}$ | $2.3 \times 10^{-6}$ | 0.0015 | $2.26 \times 10^{-7}$ | 0.0026 | 90 | 22.04 |
| 2/N2 | Exp | $2 \times 10^{-2}$ | $1.1 \times 10^{-5}$ | 0.0033 | $2.26 \times 10^{-7}$ | 0.0026 | 89 | 22.09 |
| 2/N2 | Exp | $4 \times 10^{-3}$ | $5.7 \times 10^{-5}$ | 0.0075 | $2.26 \times 10^{-7}$ | 0.0026 | 95 | 22.11 |
| 2/N2 | Exp | $8 \times 10^{-4}$ | $2.8 \times 10^{-4}$ | 0.017 | $2.26 \times 10^{-7}$ | 0.0026 | 85 | 22.47 |
| 2/N2 | Exp | $1.6 \times 10^{-4}$ | $1.4 \times 10^{-3}$ | 0.038 | $2.26 \times 10^{-7}$ | 0.0026 | 34 | 23.17 |
| 2/N2 | Exp | $3.2 \times 10^{-5}$ | $7.1 \times 10^{-3}$ | 0.084 | $2.26 \times 10^{-7}$ | 0.0026 | 9 | 23.90 |
| 2/N2 | Exp | $6.4 \times 10^{-6}$ | $3.5 \times 10^{-2}$ | 0.19 | $2.26 \times 10^{-7}$ | 0.0026 | 3 | 23.93 |
| 2/N2 | Norm | $5 \times 10^{-1}$ | $4.5 \times 10^{-7}$ | 0.00067 | $2.26 \times 10^{-7}$ | 0.0026 | 79 | 22.03 |
| 2/N2 | Norm | $1 \times 10^{-1}$ | $2.3 \times 10^{-6}$ | 0.0015 | $2.26 \times 10^{-7}$ | 0.0026 | 88 | 22.08 |
| 2/N2 | Norm | $2 \times 10^{-2}$ | $1.1 \times 10^{-5}$ | 0.0033 | $2.26 \times 10^{-7}$ | 0.0026 | 76 | 22.04 |
| 2/N2 | Norm | $4 \times 10^{-3}$ | $5.7 \times 10^{-5}$ | 0.0075 | $2.26 \times 10^{-7}$ | 0.0026 | 99 | 22.07 |
| 2/N2 | Norm | $8 \times 10^{-4}$ | $2.8 \times 10^{-4}$ | 0.017 | $2.26 \times 10^{-7}$ | 0.0026 | 94 | 22.19 |
| 2/N2 | Norm | $1.6 \times 10^{-4}$ | $1.4 \times 10^{-3}$ | 0.038 | $2.26 \times 10^{-7}$ | 0.0026 | 48 | 23.10 |
| 2/N2 | Norm | $3.2 \times 10^{-5}$ | $7.1 \times 10^{-3}$ | 0.084 | $2.26 \times 10^{-7}$ | 0.0026 | 12 | 23.82 |
| 2/N2 | Norm | $6.4 \times 10^{-6}$ | $3.5 \times 10^{-2}$ | 0.19 | $2.26 \times 10^{-7}$ | 0.0026 | 5 | 24.04 |
| 2/PB | Exp | $5 \times 10^{-1}$ | $2.1 \times 10^{-6}$ | 0.0015 | $1.06 \times 10^{-6}$ | 0.0038 | 106 | 23.39 |
| 2/PB | Exp | $1 \times 10^{-1}$ | $1.1 \times 10^{-5}$ | 0.0033 | $1.06 \times 10^{-6}$ | 0.0038 | 96 | 23.39 |
| 2/PB | Exp | $2 \times 10^{-2}$ | $5.3 \times 10^{-5}$ | 0.0073 | $1.06 \times 10^{-6}$ | 0.0038 | 122 | 23.30 |
| 2/PB | Exp | $4 \times 10^{-3}$ | $2.6 \times 10^{-4}$ | 0.016 | $1.06 \times 10^{-6}$ | 0.0038 | 109 | 23.46 |
| 2/PB | Exp | $8 \times 10^{-4}$ | $1.3 \times 10^{-3}$ | 0.036 | $1.06 \times 10^{-6}$ | 0.0038 | 37 | 25.34 |
| 2/PB | Exp | $1.6 \times 10^{-4}$ | $6.6 \times 10^{-3}$ | 0.081 | $1.06 \times 10^{-6}$ | 0.0038 | 2 | 27.92 |
| 2/PB | Exp | $3.2 \times 10^{-5}$ | $3.3 \times 10^{-2}$ | 0.18 | $1.06 \times 10^{-6}$ | 0.0038 | 0 | 29.08 |
| 2/PB | Exp | $6.4 \times 10^{-6}$ | $1.7 \times 10^{-1}$ | 0.41 | $1.06 \times 10^{-6}$ | 0.0038 | 0 | 29.45 |
| 2/PB | Norm | $5 \times 10^{-1}$ | $2.1 \times 10^{-6}$ | 0.0015 | $1.06 \times 10^{-6}$ | 0.0038 | 96 | 23.37 |
| 2/PB | Norm | $1 \times 10^{-1}$ | $1.1 \times 10^{-5}$ | 0.0033 | $1.06 \times 10^{-6}$ | 0.0038 | 110 | 23.24 |
| 2/PB | Norm | $2 \times 10^{-2}$ | $5.3 \times 10^{-5}$ | 0.0073 | $1.06 \times 10^{-6}$ | 0.0038 | 122 | 23.25 |
| 2/PB | Norm | $4 \times 10^{-3}$ | $2.6 \times 10^{-4}$ | 0.016 | $1.06 \times 10^{-6}$ | 0.0038 | 130 | 23.40 |
| 2/PB | Norm | $8 \times 10^{-4}$ | $1.3 \times 10^{-3}$ | 0.036 | $1.06 \times 10^{-6}$ | 0.0038 | 59 | 24.68 |
| 2/PB | Norm | $1.6 \times 10^{-4}$ | $6.6 \times 10^{-3}$ | 0.081 | $1.06 \times 10^{-6}$ | 0.0038 | 5 | 27.50 |
| 2/PB | Norm | $3.2 \times 10^{-5}$ | $3.3 \times 10^{-2}$ | 0.18 | $1.06 \times 10^{-6}$ | 0.0038 | 0 | 29.12 |
| 2/PB | Norm | $6.4 \times 10^{-6}$ | $1.7 \times 10^{-1}$ | 0.41 | $1.06 \times 10^{-6}$ | 0.0038 | 0 | 29.31 |

**Table S4.**
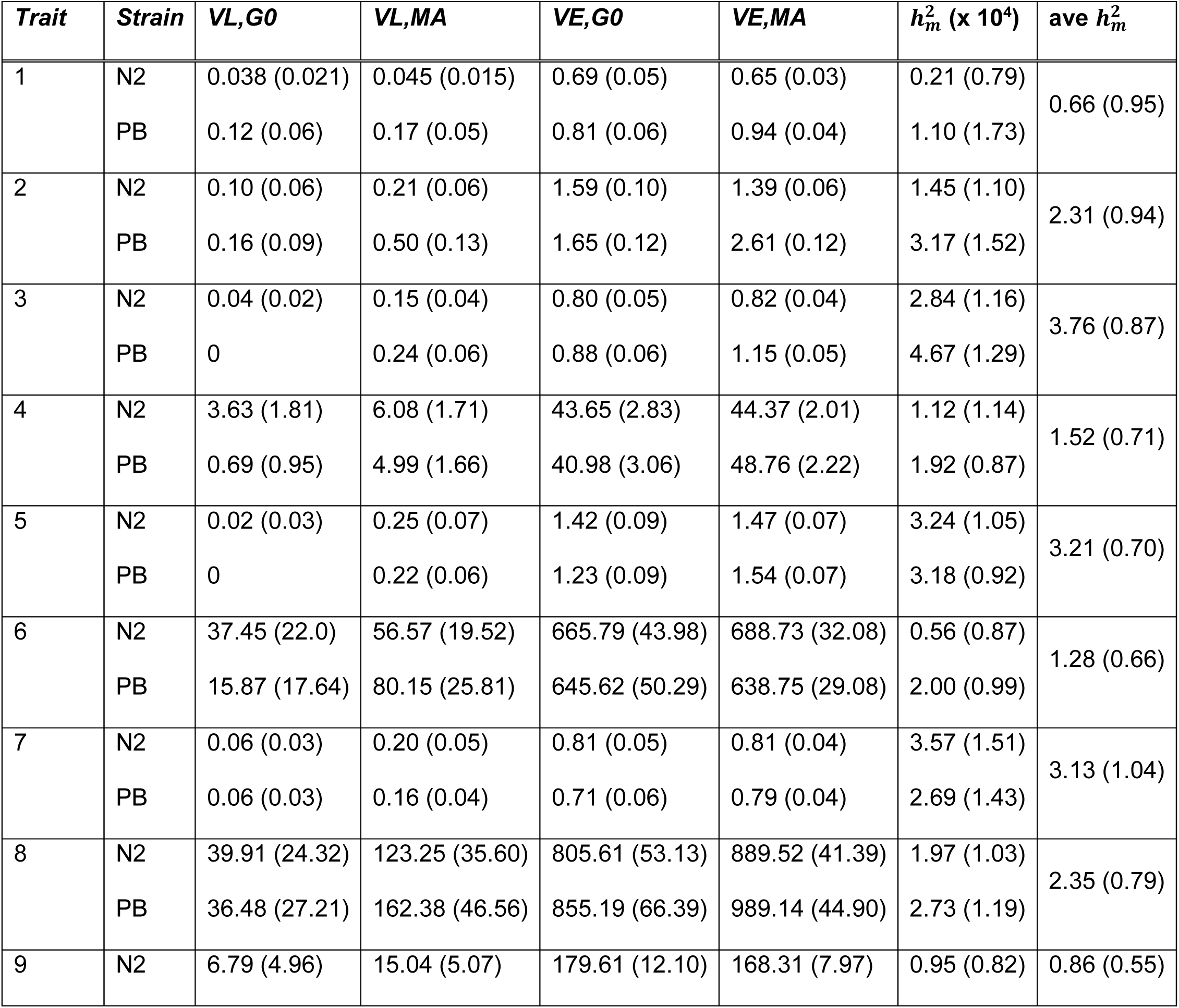
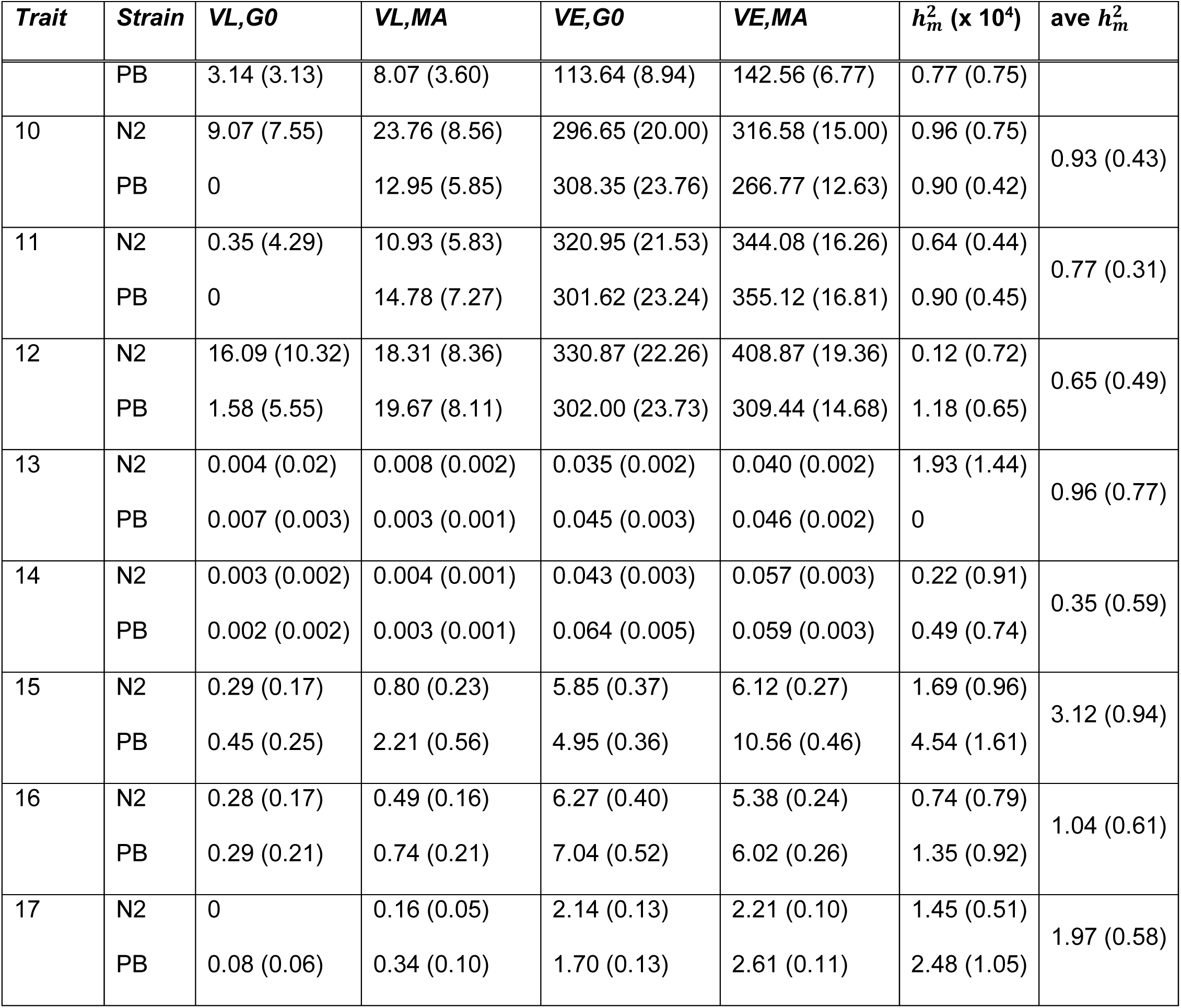
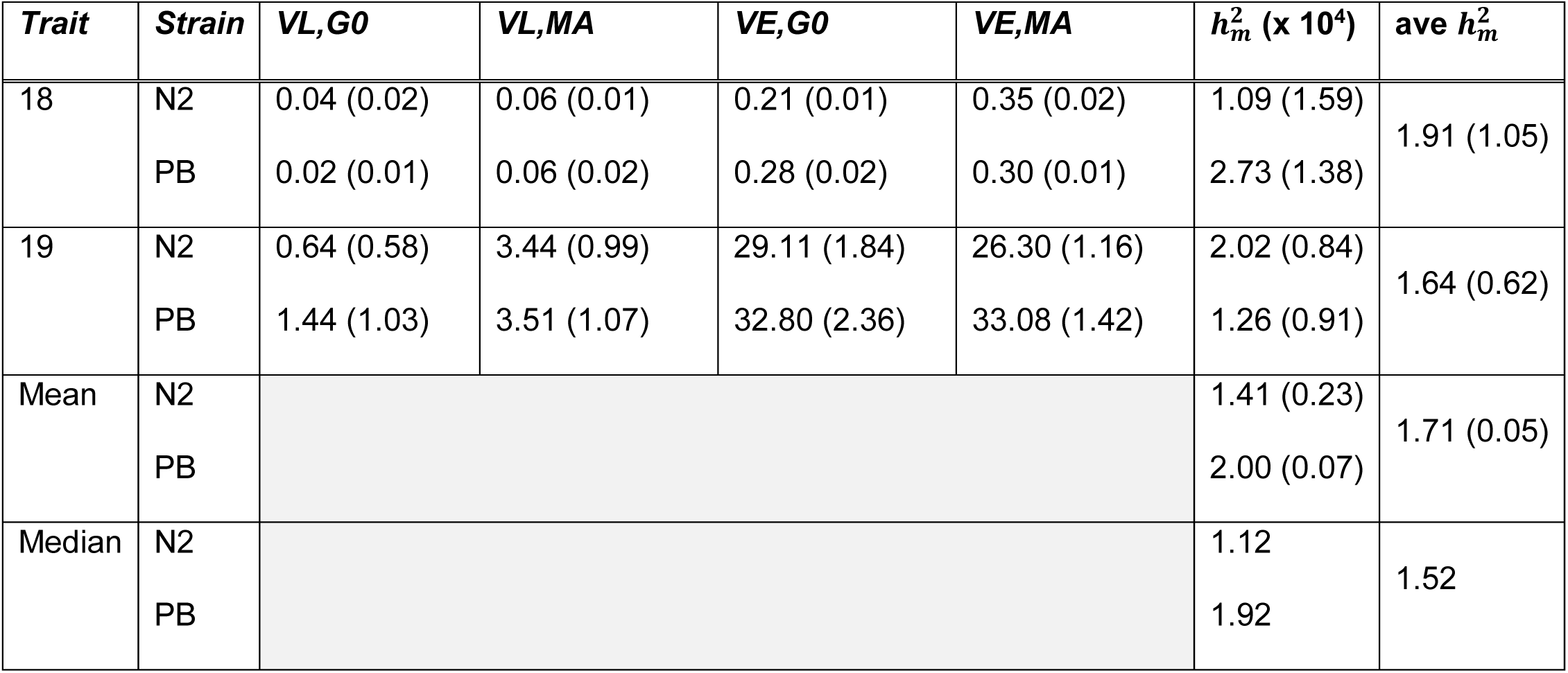
Raw variances of unstandardized traits and mutational heritabilities. Standard errors in parentheses. Column headings are: *VL,G0*, among-line variance of G0 pseudolines; *VL,MA*, among-line variance of MA lines; *VE,G0*, within-line variance of G0 pseudolines; *VE,MA*, within-line variance of MA lines; 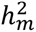, mutational heritability (× 10^4^); *ave* 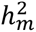, average mutational heritability of the two strains. Standard errors of 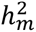 for individual traits are calculated from the square-root of the sum of the sampling variances of the G0 pseudolines and MA lines. Standard errors of the mean 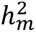, are calculated as the among-trait variance divided by the square-root of the number of traits.

**Table S5.** Variances of mean-standardized life history traits and body volume at maturity; standard errors in parentheses. All traits are from worms grown under MA conditions (on NGM agar plates at 20° C, fed on *E. coli* OP50) unless noted otherwise. Column headings are: *Trait* (units in parentheses, definitions below); *VL_MA_*, among-line variance of MA lines; *VL_G0_*, among-line variance of G0 controls; *VE_MA_*, within-line variance of MA lines; *VE_G0_*, within-line variance of G0 controls; *VM*, mutational variance; 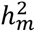, mutational heritability; VG, genetic variance. Trait abbreviations are: W20, lifetime reproduction weighted by survival; *W25*, lifetime reproduction weighted by survival at 25° C; *W_SORT_*, lifetime reproduction of worms grown individually in liquid media in microplates; *Surv*, proportion of embryos surviving to 72 hrs; *LT50MA*, median lifespan under MA conditions; *LT50Pa*, median lifespan of worms exposed to the pathogenic bacteria *Pseudomonas aeruginosa*; *Size*, body volume at maturity. VG for traits marked with an asterisk is not estimated from the same set of wild isolates included in this study. Experimental details are reported in Etienne et al. 2015.

| *Trait* | $VL_{MA} (x 10^3)$ | $VL_{G0} (x 10^3)$ | $VE_{MA} (x 10^2)$ | $VE_{G0} (x 10^2)$ | $VM (x 10^5)$ | $h_m^2 (x 10^3)$ | $VG (x 10^3)$ | $t_P$ |
| --- | --- | --- | --- | --- | --- | --- | --- | --- |
| *W20\** (# offspr) |  |  |  |  | 15.49 | 1.57 | 13.7 | 88.4 |
| *W25* (# offspr) | 99.11 (39.03) | 9.71 (33.96) | 34.63 (3.81) | 48.01 (6.22) | 20.32 (11.76) | 0.49 |  |  |
| *W<sub>Sort</sub>* (# offspr) | 55.57 (12.74) | 3.05 (1.96) | 19.78 (2.12) | 14.90 (2.29) | 10.50 (2.58) | 4.64 | 17.5 | 166.7 |
| *Surv* (Pct) | 16.72 (10.32) | 0 | 16.58 (1.50) | 12.62 (1.38) | 3.39 (2.64) | 0.23 |  |  |
| *LT50MA* (hrs) | 33.37 (9.43) | 3.91 (4.49) | 2.52 (0.40) | 1.42 (4.62) | 5.72 (2.10) | 2.91 |  |  |
| *LT50Pa* (hrs) | 8.97 (3.26) | 0 | 1.62 (1.56) | 1.45 (2.22) | 1.79 (0.65) | 1.17 | 6.0 | 335.2 |
| *Size\** (mm<sup>3</sup>) | 54.15 (15.53) | 1.08 (3.66) | 36.93 (7.32) | 37.63 (7.22) | 13.35 (3.19) | 3.58 | 2.3 | 17.2 |
| *Mean / Median* | 44.6 / 43.8 | 3.0 / 2.1 | 18.7 / 18.2 | 19.3 / 13.8 | 9.2 / 8.1 | 2.2 / 2.0 | 9.9 / 9.9 |  |

